# RAD52 and RPA act in a concert promoting inverse RNA strand exchange

**DOI:** 10.1101/2025.07.22.666130

**Authors:** Sarah F. DiDomenico, Hoang H. Dinh, Matthew J. Rossi, Antoine Baudin, Simranjeet S. Sekhon, Walter J. Chazin, David S. Libich, Alexander V. Mazin

## Abstract

Recent studies in eukaryotes have revealed an important role of RNA in DNA repair and identified the RAD52 protein as a central player in RNA-dependent repair of DNA. In vitro, RAD52 promotes inverse RNA strand exchange between dsDNA and homologous RNA. This reaction is strongly stimulated by the RAD52 partner, replication protein A (RPA). Here, using NMR and biochemical methods we investigated the mechanism of this stimulation. We identified two RPA-binding sites in the unstructured RAD52 C-terminal domain (CTD), which mediate interaction with RPA70 and RPA32 subunits.

These interactions are critical for stimulation of inverse RNA strand exchange. Furthermore, we showed that stimulation of inverse RNA strand exchange requires formation of an RPA-RNA complex that strengthens the RPA-RAD52 interaction and serves to deliver RNA to the RAD52-dsDNA complex for strand exchange. These results elucidate the mechanism of novel inverse RNA strand exchange activity of RAD52 and the role of RAD52-RPA interaction in RNA-dependent DNA repair.

## INTRODUCTION

DNA double-strand breaks (DSBs), the most deleterious DNA lesions, cause genome instability which lead to cancer and other genetic diseases (Hanahan and Weinberg 2011, Zhao, Wiese et al. 2019). The homologous recombination (HR) pathway promotes DSB repair with high fidelity as opposed to error-prone DSB repair pathways like Non-Homology End-Joining (Kowalczykowski 2015).

In yeast, the Rad52 protein plays a critical role in all HR events (Game and Mortimer 1974, Sung 1997, Krogh and Symington 2004, Mortensen, Lisby et al. 2009). In mammals, RAD52 knockouts do not show strong deficiencies in DNA repair or HR (Rijkers, Van Den Ouweland et al. 1998). Instead, RAD52 mutations cause synthetic lethality in HR-deficient cancer cells that carry mutations in BRCA1, BRCA2 and several related HR genes (Feng, Scott et al. 2011, Jalan, Olsen et al. 2019), highlighting the importance of RAD52 in back-up DNA repair mechanisms (Rossi, DiDomenico et al. 2021). Recent data indicate that RAD52 plays an important role in RNA-dependent DNA repair (Keskin, Shen et al. 2014, Wei, Nakajima et al. 2015, Mazina, Keskin et al. 2017, Welty, Teng et al. 2018, Yasuhara, Kato et al. 2018). In these and other studies, RNA was shown to stimulate HR by serving as a template, primer, and/or signal molecule during DNA repair or recombination (Storici, Bebenek et al. 2007, Meers, Keskin et al. 2020, Tan, Duan et al. 2020, Chandramouly, Zhao et al. 2021, Jalan, Brambati et al. 2025) Human RAD52 is a 47 kDa protein of 418 amino acid (aa) residues comprised of two equally sized N-terminal (NTD, 1-209) and C-terminal (CTD, 210-418) domains (Hanamshet, Mazina et al. 2016, Balboni, Rinaldi et al. 2023). The highly conserved RAD52 NTD contains two DNA binding sites and an oligomerization domain responsible for formation of a RAD52 undecameric ring (Kagawa, Kurumizaka et al. 2002, Singleton, Wentzell et al. 2002, Kinoshita, Takizawa et al. 2023, Balboni, Marotta et al. 2024, Liang, Greenhough et al. 2024, Honda, Razzaghi et al. 2025) and is sufficient for all RAD52 DNA and RNA pairing activities (Sugiyama, New et al. 1998, Kagawa, Kurumizaka et al. 2001, Lloyd, McGrew et al. 2005, Kagawa, Kagawa et al. 2008, Grimme, Honda et al. 2010, Mazina, Keskin et al. 2017). The structure of the less evolutionary conserved CTD was not resolved likely due to its predicted intrinsically disordered nature (Balboni, Marotta et al. 2024, Struble, Lovelace et al. 2024). The CTD contains the nuclear localization signal and sites for binding of RAD52 protein partners including RAD51 recombinase and replication protein A (RPA), a major eukaryotic ssDNA binding protein (Park, Ludwig et al. 1996, Koike, Yutoku et al. 2013, Ma, Kwon et al. 2017). However, the role of these interactions and the overall function of the RAD52 CTD in DNA repair and genome stability in mammalian cells remains to be elucidated.

In vitro, RAD52 promotes annealing between complementary ssDNA or RNA molecules (Mortensen, Bendixen et al. 1996, Sugiyama, New et al. 1998, Grimme, Honda et al. 2010, Keskin, Shen et al. 2014). The RAD52-ssDNA complex can also promote DNA strand exchange producing joint molecules or D-loops (Kagawa, Kurumizaka et al. 2001). More recently, we showed that RAD52 promotes inverse RNA strand exchange, the reaction in which RAD52 forms a complex with dsDNA and promotes strand exchange with homologous RNA to form an RNA:DNA heteroduplex. This RAD52 activity was demonstrated to play a role in RNA-templated DNA repair in yeast (Mazina, Keskin et al. 2017).

Previously, we showed that the inverse RNA strand exchange activity of RAD52 is strongly stimulated by RPA and that this stimulation depends on the RAD52 CTD (Mazina, Keskin et al. 2017). Here, we aimed to characterize the mechanism of this stimulation. The RPA is a heterotrimer comprised of three subunits, RPA70, RPA32, and RPA14. Previously, it was shown that a region of RPA32 (172-270), RPA32C, interacts with the RAD52 CTD region (257-274) (Park, Ludwig et al. 1996, Mer, Bochkarev et al. 2000). First, we utilized solution NMR spectroscopy to refine the RPA binding region in the RAD52 CTD. Our data not only confirmed that RPA binds to the previously identified RAD52 CTD site (247-280) but revealed a novel RPA binding site between residues 308-335. We showed that these two sites, defined as the primary and secondary, bind separately to RPA RPA32C domain and the RPA70 N-terminal domain (1-120) (RPA70N) that is another major RPA interaction module (Bochkareva, Kaustov et al. 2005, Xu, Vaithiyalingam et al. 2008). We demonstrated that both RPA binding sites on RAD52 CTD contribute to stimulation of inverse RNA strand exchange, with the primary site playing a major role. Furthermore, we found that formation of an RPA-RNA complex is essential for stimulation of inverse RNA strand exchange in two ways: by strengthening the RAD52-RPA interaction and by delivering RNA to RAD52. These results help to clarify the mechanism of stimulation of inverse RNA strand exchange activity of RAD52 by RPA.

## RESULTS

### RAD52-RPA interaction is enhanced by ssDNA or RNA

We previously showed that RAD52 performs inverse RNA strand exchange *in vitro* by forming a complex with dsDNA and performing strand exchange with homologous RNA to produce an RNA:DNA hybrid. The reaction is strongly stimulated by RPA. The RAD52 NTD promotes inverse RNA strand exchange efficiently, however its activity is not stimulated by RPA (Mazina, Keskin et al. 2017) indicating that RPA binding to the CTD is essential for the stimulation. Here, we wanted to understand the mechanism of this stimulation.

RAD52 interacts with free RPA (Park, Ludwig et al. 1996, Mer, Bochkarev et al. 2000, Jackson, Dhar et al. 2002) or RPA-ssDNA complexes (Ma, Kwon et al. 2017), and since RPA binds RNA (Mazina, Somarowthu et al. 2020), we endeavored to determine if RAD52 also interacts with RPA-RNA complexes. We expressed and purified from *E. coli* glutathione S-transferase (GST)-tagged RAD52 CTD that contains the RPA binding region to test for RPA binding. Using a pull-down assay, we observed the interaction of the GST-RAD52-CTD with RPA (Figure 1A, lane 10). The interaction was enhanced by ssDNA or RNA added to RPA in equimolar concentrations (Figure 1A, lanes 4 and 7).

**Figure 1.**
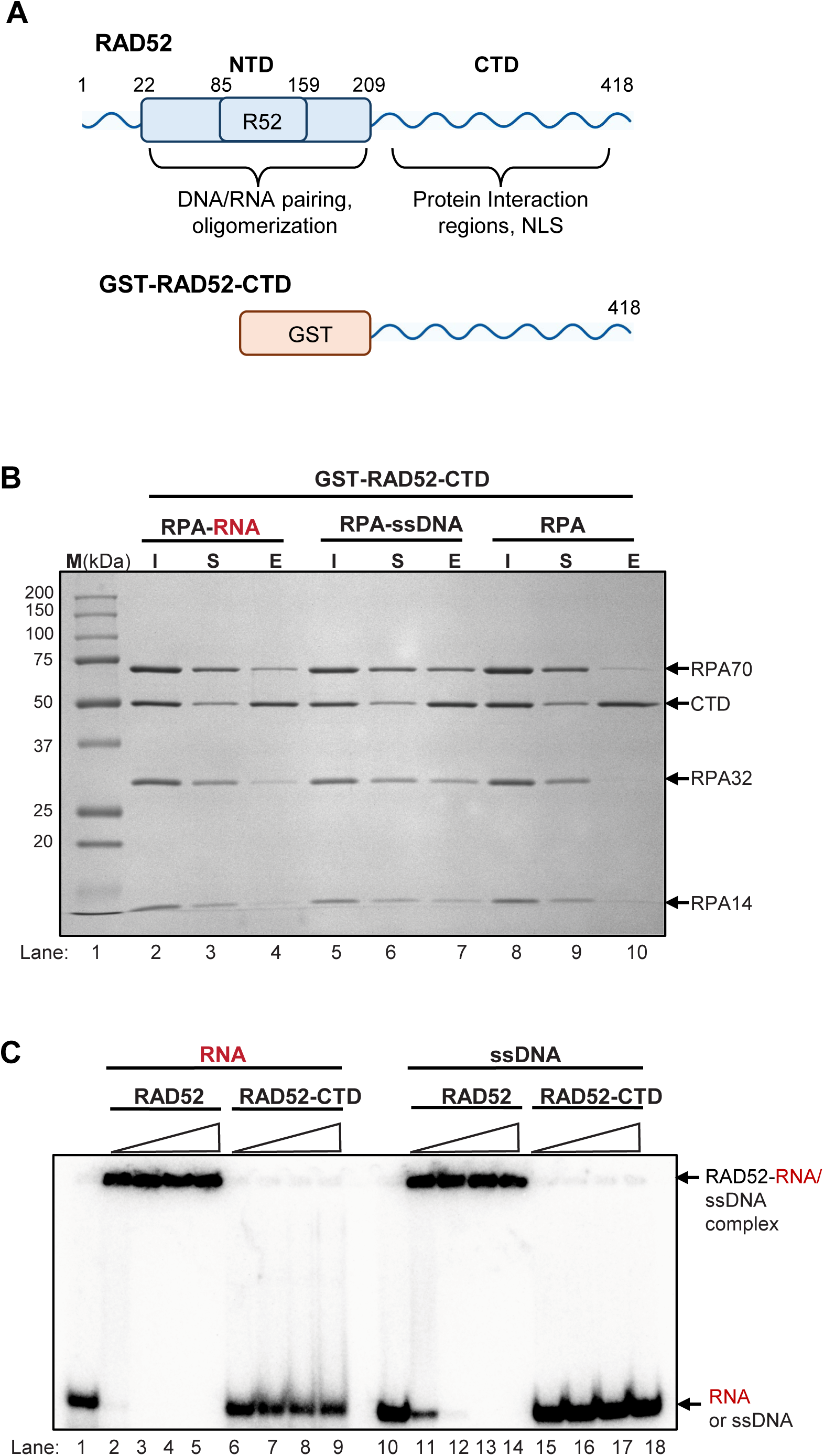
RAD52-RPA Interaction is strengthened by RNA or ssDNA. (**A**) The schematic of the RAD52 full length (1-418) and GST-tagged RAD52 CTD (217-418). The wavy lines indicate predicted protein disordered regions. (**B**) The interaction of GST-RAD52-CTD (6 µM) with RPA (6 µM); the RPA complex (6 µM) with RNA (no.517, 63-mer, 6 µM); or RPA complex (6 µM) with ssDNA (no.211, 48-mer, 6 µM) was assessed by affinity pulldown. Proteins were analyzed by gel electrophoresis in 12% SDS-PAGE with Coomassie blue staining. Lanes are input (I), supernatant (S), and elution (E). (**C**) RAD52 full length but not RAD52-CTD binds RNA or ssDNA. Full length RAD52 or RAD52-CTD (0.5, 1, 1.5, or 2 µM) was incubated with ^32^P-labeled ssDNA (no. 1, 63 nM) or RNA (no. 517, 63 nM). Complex formation was analyzed by electrophoresis in 8% polyacrylamide gels. Experiments were repeated at least 3 times. Shown is a representative gel image.

Since the RAD52 CTD does not bind either ssDNA (Jackson, Dhar et al. 2002) or RNA (Figure 1B), these results indicated that the stimulation was due to the enhancement of protein:protein interactions between RAD52 CTD and RPA. This is consistent with the report that ssDNA binding causes a change in RPA domain configuration (Brosey, Chagot et al. 2009, Fan and Pavletich 2012), which may facilitate these interactions.

### RAD52 has two distinct RPA binding sites

Next, using NMR we wanted to identify RNA binding site(s) in the intrinsically disordered RAD52 CTD (Suppl. Figure S1A). The^1^H-^15^N heteronuclear single quantum coherence (HSQC) spectra of RAD52 CTD (217-418) revealed narrow ^1^H chemical shift dispersion and overlapped peaks, characteristic of intrinsic disorder (Suppl. Figure S1B). Using standard approaches (Johnson, Xu et al. 2022, Baudin, Dinh et al. 2025) we unambiguously assigned 81% of non-proline backbone resonances (Suppl. Figure S1B). Secondary structure propensity (SSP) calculations (Marsh, Singh et al. 2006) using chemical shift data from protein backbone atoms (^1^H_N_, ^15^N, ^13^C⍺, ^13^Cβ, and ^13^C’) revealed that the RAD52 CTD fragment is largely disordered, except residues 248-278 which are predicted to adopt an ⍺-helical conformation (Suppl. Figure S2). We measured ^15^N *T*_1_ and *T*_2_ relaxation, and ^1^H-^15^N heteronuclear NOEs (hetNOEs) to assess RAD52 CTD fast-timescale dynamics. Average *R*_1_ and *R*_2_ rates were 1.5 s^-1^ and 4.4 s^-1^, respectively, with residues 248-278 showing elevated *R*_2_ (∼8 s^-1^), consistent with stable helices. The average hetNOE was 0.15, with higher values (∼0.5) for residues 248-278, indicating restrained motion typical of secondary structure (Suppl. Figure S2). These data suggest that while most of the RAD52 CTD is disordered, residues 248-278 likely form a stable ⍺-helix.

In early studies, an RPA binding site was identified spanning residues 247-280 of the RAD52 CTD, within the region of our observations of ⍺-helical structure (Figure 2A) (Park, Ludwig et al. 1996). To confirm that RAD52 CTD residues 248-278 are involved in RPA binding, we mapped RPA’s interaction with the RAD52 CTD using chemical shift perturbation (CSP) analysis. We conducted NMR CSP and line broadening analysis using ^15^N-labeled RAD52-CTD (217-418) titrated with unlabeled (NMR-invisible) full-length RPA trimer (RPA70:RPA14:RPA32) (Figure 2B). We observed only small CSPs (< 0.10 ppm) for a handful of residues (e.g. 279, 315) in the 247-280 and 308-335 regions (Figures 2A,C; Suppl. Figure S3A). More significantly, the intensities of all RAD52-CTD peaks were uniformly reduced ∼65%, an observation consistent with the 22.4 kDa RAD52 CTD interacting with the much larger 111.2 kDa RPA hetrotrimer. For residues 247-280 and 308-335, the observed broadening was more severe than the average with many peaks within these regions, such as R260, Q261, and V317, completely disappearing from the ^1^H,^15^N-HSQC spectrum (Figure 2C; Suppl. Figure S3A). This differential broadening likely arises from lifetime line broadening effects due to direct contact between these two RAD52-CTD regions and the RPA heterotrimer, however, we cannot rule out contributions from conformational exchange within these regions (Libich, Fawzi et al. 2013).

**Figure 2.**
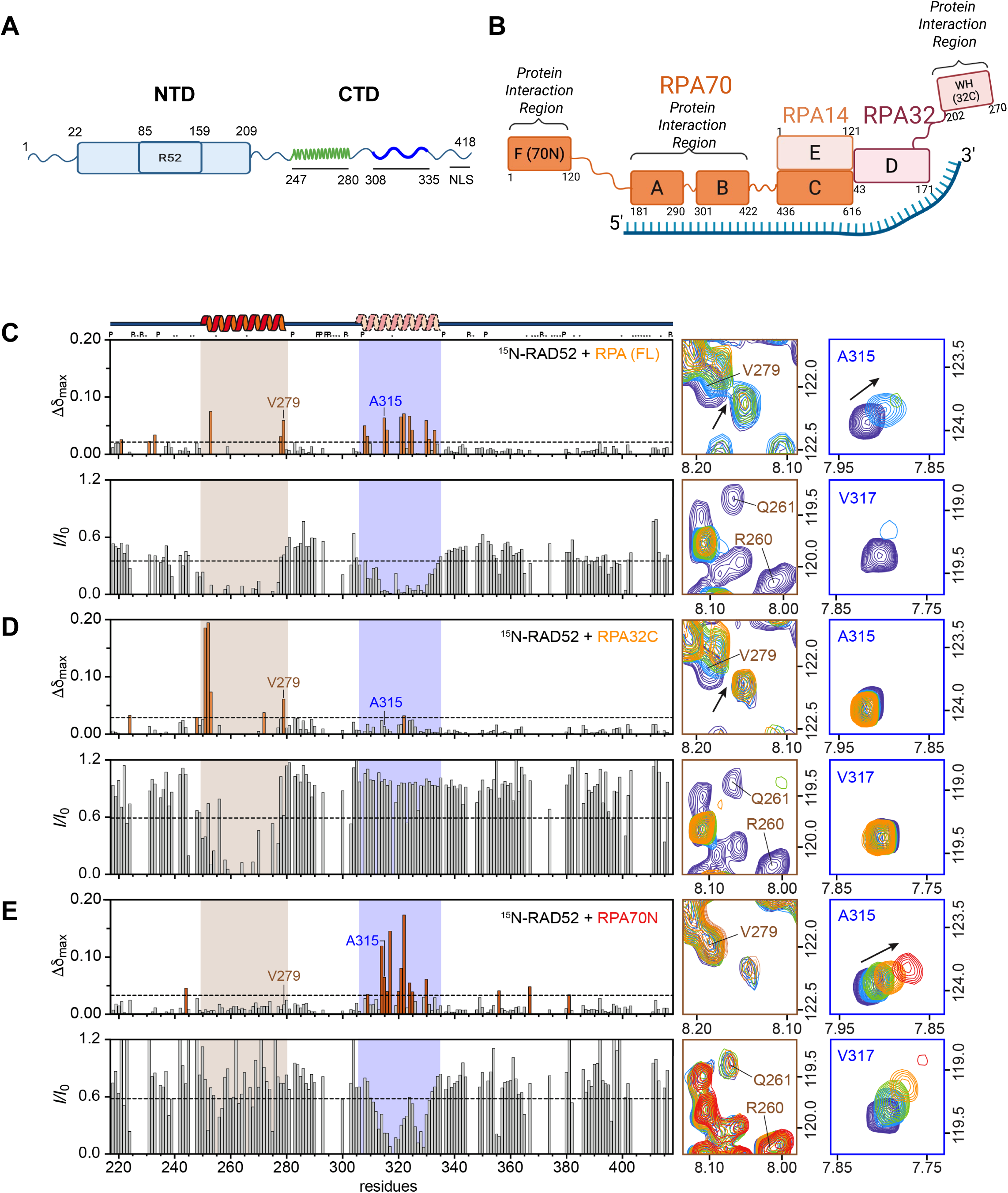
RAD52 has two distinct RPA Binding Sites. (**A**) The schematic of RAD52 showing the ordered NTD from residues 1-209 and the disordered CTD from 210-418. The oligomerization site (R52) is between residues 85-159. The nuclear localization sequence (NLS) is between residues 405-414. (**B**) Schematic of the RPA heterotrimer comprised of RPA70 (orange), RPA32 (red), and RPA14 (light orange) subunits. The domains of each subunit that interact with ssDNA are shown bound to ssDNA (blue). RAD52-CTD (217-418) was used in NMR experiments to map the interaction region(s) with full length RPA heterotrimer, RPA32C (202-270), or RPA70N (1-120) domains. (**C-E**) Plots showing the maximum change in chemical shift (Δδ_max_ (ppm)) (top) and average change in peak intensity (*I*/*I*_0_) (bottom) of the assigned backbone amides of RAD52-CTD when titrated with (**C**) RPA heterotrimer, (**D**) RPA32C or (**E**) RPA70N domains.

### The primary and secondary CTD sites interact with RPA32C and 70N

Next, we wanted to identify the specific RPA domains interacting with the primary and secondary RAD52 CTD sites (Figure 2A). It was reported that the C-terminal region of the RPA32 subunit (RPA32C) (172-270) (Figure 2B) interacts with the RAD52 CTD between residues 257-274 (overlapping with the primary site) (Park, Ludwig et al. 1996, Mer, Bochkarev et al. 2000). Our titrations of the RAD52 CTD with full-length RPA clearly identified a second interaction site (308-335) that is distinct from the RPA32C binding site (Figure 2C). In addition to the RPA32C subunit, the RPA70 N-terminal domain subunit (RPA70N) (1-120) has been described as a major protein-protein interaction module of RPA (Figure 2B) (Bochkareva, Kaustov et al. 2005, Xu, Vaithiyalingam et al. 2008, Prakash and Borgstahl 2012).

To identify which subunit(s) of the RPA heterotrimer interact with the RAD52 CTD, we titrated ^15^N-labeled RAD52-CTD with unlabeled RPA70N or RPA32C domain constructs. With the addition of RPA32C, only residues within the primary RPA binding site (247-280) showed concentration dependent CSPs and differential line-broadening, while residues within the secondary site (308-335) remained unperturbed (Figure 2D; Suppl. Figure 3B). Additionally, a uniform reduction in RAD52 CTD peak intensities was not observed since the RPA32C subunit is only 7.6 kDa. These observations are consistent with previous reports of RPA32C interacting with residues 257-274 of the RAD52 CTD (Park, Ludwig et al. 1996, Mer, Bochkarev et al. 2000) and further support a separate region of RPA interacting with the secondary RPA binding site (308-335) in the RAD52 CTD. Next, we titrated ^15^N RAD52 CTD with unlabeled RPA70N and observed CSPs > 0.10 ppm for L314, V317, and I322 residues, and no appreciable perturbations within the helical 247-280 region (Figure 2E; Suppl. Figure S3C). An observed uniform reduction in intensity of ∼10-20% for all residues is consistent with RPA70N being larger (13.2 kDa) than RPA32C, and a greater reduction in intensity for residues RAD52 CTD 308-335 is indicative of a direct interaction with RPA70N (Figure 2E). Taken together, our NMR data suggest that two distinct regions on RAD52 CTD are responsible for its interaction with the RPA heterotrimer, a primarily helical region (247-280) that binds to RPA32C, and the secondary site (308-335) that binds to RPA70N (Figure 2A).

### Identification of the RAD52 residues important for interaction with RPA

To further characterize the role of the two sites identified by NMR analysis in the RAD52-RPA interaction, we constructed and purified GST-tagged RAD52 CTD (217-418) with point mutations (Figure 3A). Consistent with previous reports, the primary site contains the RQK (R260/Q261/K262) motif that mediates interactions with RPA32C (Mer, Bochkarev et al. 2000, Grimme, Honda et al. 2010). Here, we generated charge swapping mutations R260Q/Q261E/K262M (RAD52 CTD^QEM^) to maximize disruption of the RAD52 CTD interaction with RPA32C. In the secondary site, we mutated a patch of hydrophobic residues, L314A/V317N/L321Q/I322Q/L325A/W332A (RAD52 CTD^ANQ^) (Figure 3A). These RAD52 CTD protein constructs were tested in pull down assays for interaction with an RPA-RNA complex. We found that the QEM mutation disrupted the interaction between RAD52 CTD and the RPA-RNA complex (Figure 3B) in accordance with the previously reported role of the RQK motif in RAD52-RPA interaction (Grimme, Honda et al. 2010). However, the ANQ mutation only slightly decreased but did not abolish this interaction (Figure 3B). Taken together, these data suggest the primary site (247-280) plays a major role in RAD52-RPA interactions with a lesser contribution from the secondary (308-335) site.

**Figure 3.**
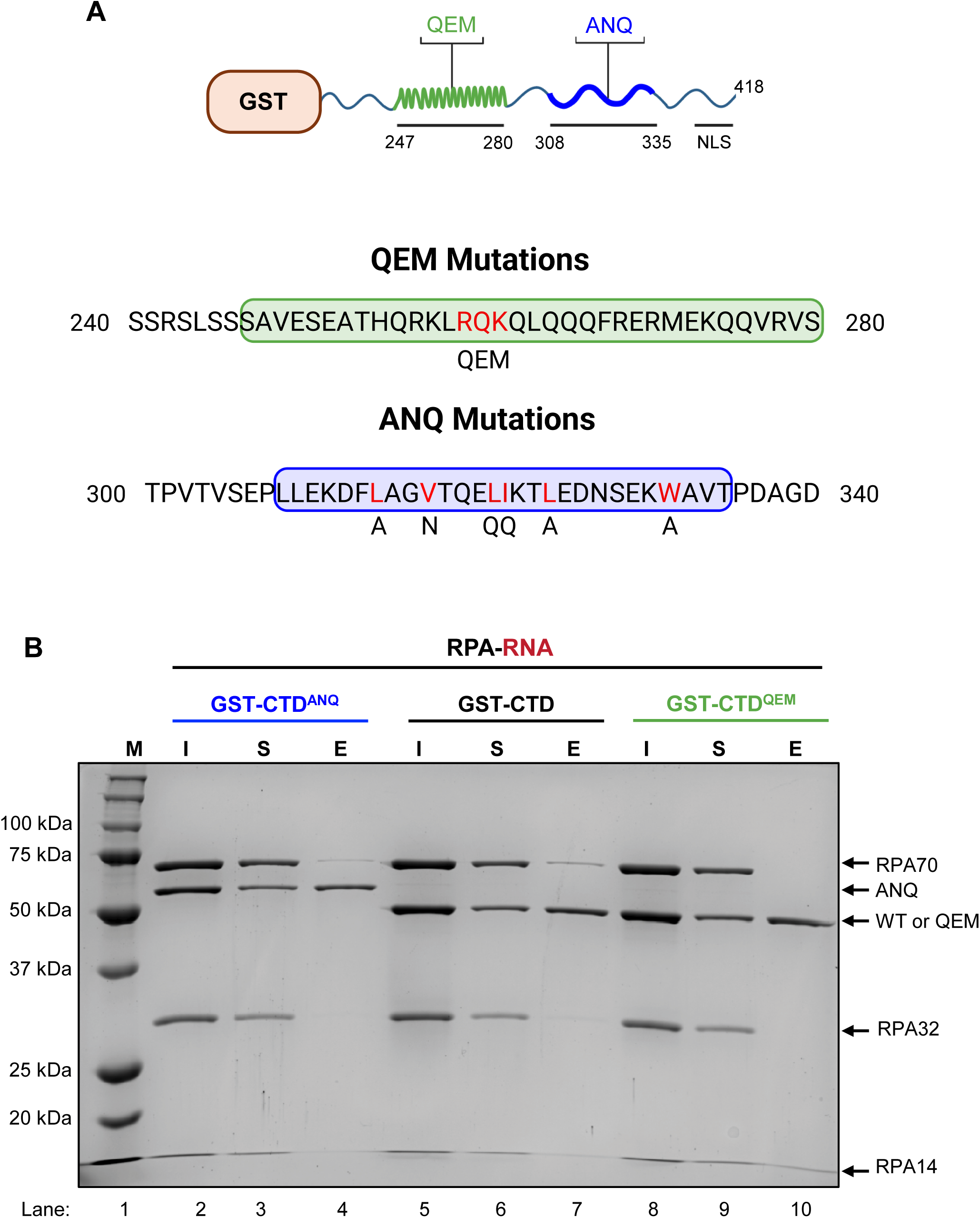
The RAD52 CTD primary site plays a dominant role in RPA binding. (**A**) Schematic representation of the GST-tagged RAD52 CTD (217-418) wild type, R260Q/Q261E/K262M (QEM), and L314A/V317N/L321Q/I322Q/L325A/W332A (ANQ) mutants. The primary (247-280, green) and secondary (308-335, blue) RPA-binding sites are highlighted and mutated residues (black) are indicated below wild-type (red) residues. (**B**) The interaction between RPA and wild-type or mutant RAD52 proteins was assessed by affinity pulldown. GST-tagged RAD52 CTD, RAD52 CTD^QEM^, or RAD52 CTD^ANQ^ (6 µM) was immobilized on glutathione sepharose (Cytiva) and incubated with an RPA-RNA (6 µM) complex. Proteins were analyzed by electrophoresis in 12% SDS-PAGE with Coomassie blue staining. Marker (M), input (I), supernatant (S), elution (E). The experiments were repeated at least 3 times.

### Both RAD52 CTD cites play a role in stimulation of inverse RNA strand exchange by RPA

To determine which specific RAD52 CTD interaction sites and residues are required for stimulation of inverse RNA strand-exchange activity by RPA, we purified full-length RAD52 with QEM (RAD52^QEM^), ANQ (RAD52^ANQ^), both QEM and ANQ (RAD52^QEM+ANQ^), and a CTD truncation (RAD52^Δ251-418^) devoid of both primary and secondary RPA binding sites. We tested these mutants along with RAD52 wildtype in inverse RNA strand exchange in the presence or absence of RPA (Figure 4A). As expected, RAD52 promoted inverse RNA strand exchange and was stimulated by RPA (Figure 4B,D; Suppl. Figure S4A). The RAD52^Δ251-418^, RAD52^QEM^, RAD52^ANQ^ and RAD52^QEM+ANQ^ promoted the reaction as efficiently as the wildtype protein (Figures 4C,D; Suppl. Figures S4B-E). However, the activity of the RAD52^Δ251-418^, RAD52^QEM^, and RAD52^QEM+ANQ^ mutants deficient in RPA32C binding were not stimulated by RPA (Figures 4C-D; Suppl. Figures S4B,C,E), indicating the primary site and, particularly the RQK residues, are required for stimulation of inverse RNA strand exchange.

**Figure 4.**
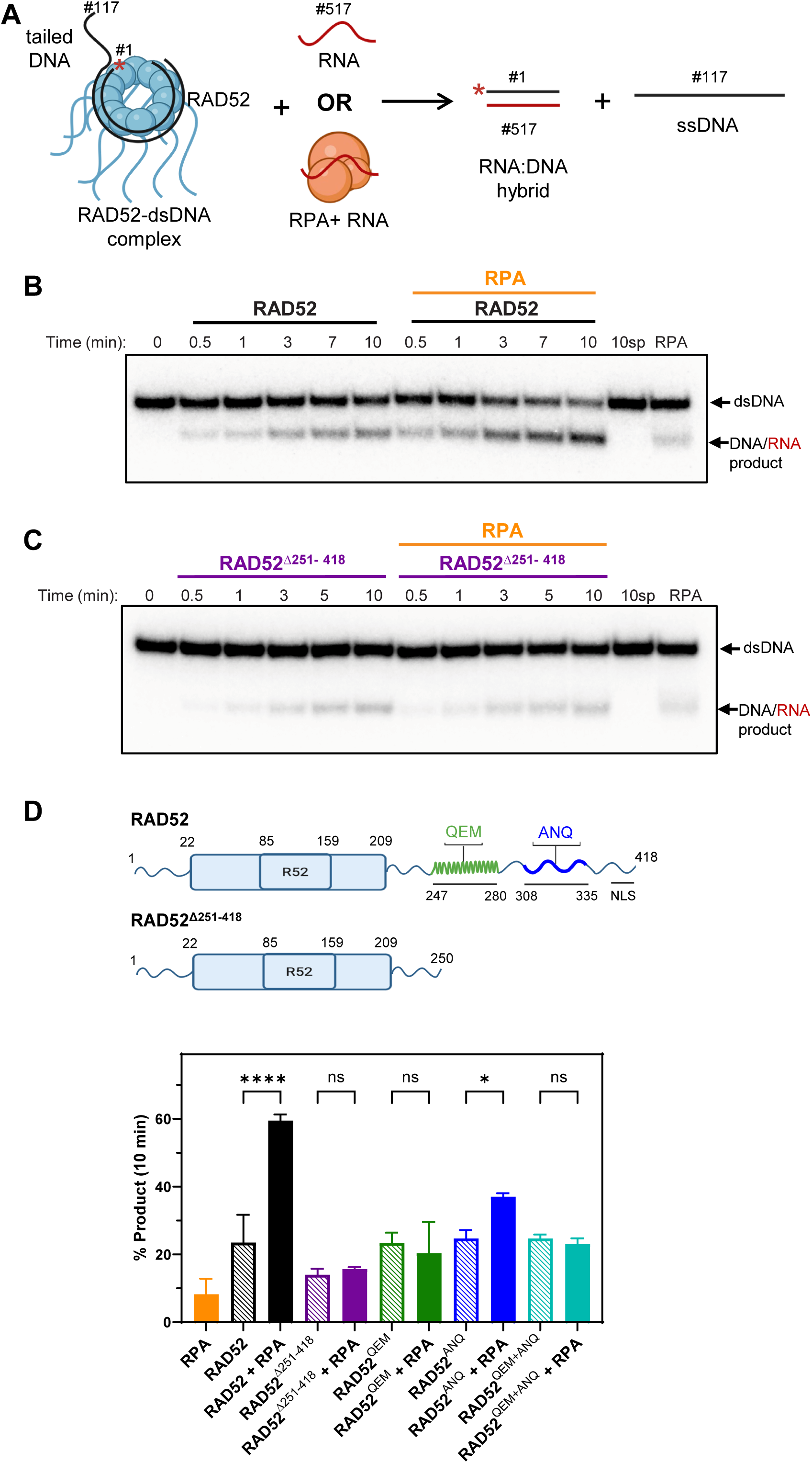
Both the primary and secondary RAD52 CTD sites are important for stimulation of inverse RNA strand exchange by RPA. (**A**) The scheme of inverse RNA strand exchange. Asterisk represents ^32^P-label. Oligonucleotide sequences are shown in Table S1. Wild-type or mutant RAD52 in concentrations indicated below were incubated with the 3’ tailed DNA (no. 117/ no. 1; 68.6 nM). The reactions were initiated by the addition of RNA (no. 517; 686 nM) alone or in complex with RPA (750 nM). (**B-C**) The representative 8% polyacrylamide gels of inverse RNA strand exchange promoted by (**B**) wildtype RAD52 (1.1 µM) or (**C**) RAD52^Δ251-418^ (2 µM). (**D**) Quantification of product formation after 10 min of inverse RNA strand exchange promoted by RAD52, RAD52^Δ251-418^ (2 µM), RAD52^QEM^ (2 µM) (mutated primary site), RAD52^ANQ^ (1.7 µM) (mutated secondary site), and RAD52^QEM+ANQ^ (1.5 µM) (double mutant) with and without RPA. The protein maps are shown at the top. The experiments were repeated at least 3 times; error bars indicate SD.

Interestingly, the secondary site RAD52^ANQ^ mutant also decreased the level of stimulation by RPA, although to a lesser extent than the primary site mutants (Figure 4D; Suppl. Figure S4D). This suggests that while the RQK motif plays a dominant role in stimulation, transient interactions with the secondary site still contribute to the overall increase of inverse RNA strand exchange activity by RPA. Together, these results suggest that both RPA interaction sites in the RAD52 CTD are important for stimulation of inverse RNA strand exchange.

### RPA32C and RPA70N domains do not stimulate inverse RNA strand exchange

Since the interaction of RPA32C and RPA70N with the RAD52 CTD contributes to stimulation of inverse RNA strand exchange activity, we examined if interaction with the isolated RPA domains was sufficient to stimulate this reaction (Figure 5A). We found that in the range of tested concentrations neither RPA32C (0-2 μM), RPA70N (0-2 μM), or a mixture of RPA32C and RPA70N (0-5.5 μM) stimulated inverse RNA strand exchange activity in contrast to full-length RPA (Figures 5B-D). These results indicate that the action of the entire RPA heterotrimer is needed since neither RPA32C nor RPA70N domain were sufficient for inverse RNA strand exchange stimulation.

**Figure 5.**
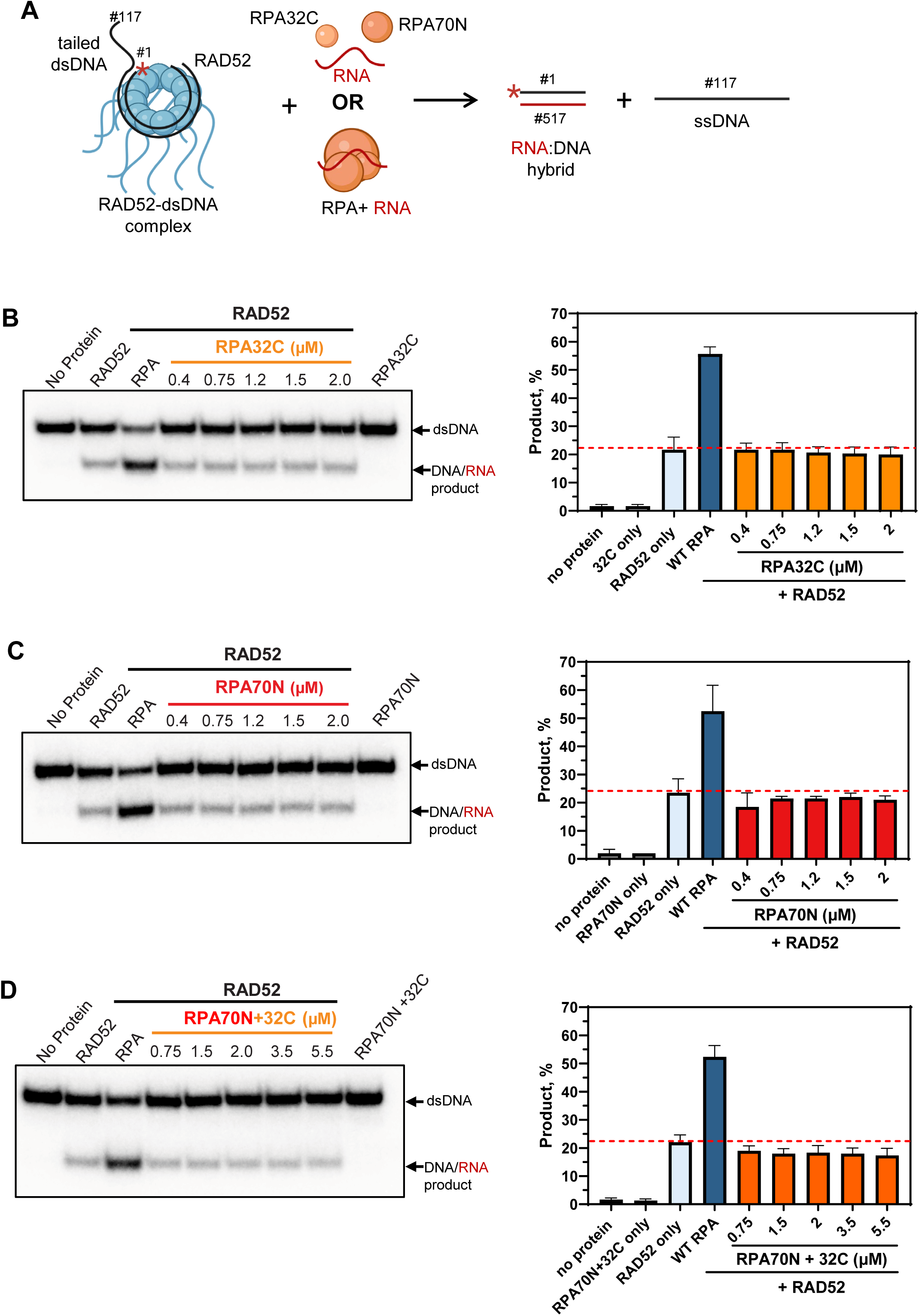
Isolated RPA32C and RPA70N domains do not stimulate RAD52 inverse RNA strand exchange. (**A**) The scheme of inverse RNA strand exchange. The reaction conditions are the same as in Figure 4 except (**B**) RPA32C, (**C**) RPA70N (0, 0.4, 0.75, 1.2, 1.5, 2 µM), or (**D**) RPA70N and RPA32C 1:1 (0, 0.75, 1.5, 2, 3.5, 5.5 µM, of each domain) were used. For RPA32C, RPA70N or RPA70N+RPA32C reactions in the absence of RAD52, the highest protein concentration was used, 2 µM or 5.5 µM, respectively. Products were analyzed by 8% polyacrylamide gel electrophoresis (left panels) and quantified (right panels). The experiments were repeated at least 3 times; error bars indicate SD.

### RPA-RNA complex formation is required for stimulation of inverse RNA strand exchange

Here, we wanted to investigate the RPA attributes that in addition to RPA32C and RPA70N binding are important for stimulation of the RAD52 inverse RNA strand exchange activity. We showed above that RNA binding to RPA strengthens its interaction with RAD52 (Figure 1A). This enhanced interaction may contribute to two potential mechanisms of stimulation. In the first, RPA-RNA binding to RAD52 may activate strand exchange activity by inducing conformational changes in RAD52. In the second, RPA acts as a shuttle by delivering homologous RNA to RAD52. The first mechanism implies that an RPA-RNA complex of RPA activates RAD52 for inverse RNA strand exchange that proceeds through interaction of RAD52 with free (non-bound to RPA) RNA. Whereas the second mechanism suggests that the reaction does not require the presence of free RNA and depends on the RPA-RNA complex.

We tested whether free (RPA unbound) RNA is required for RPA stimulation of inverse RNA strand exchange. For this, in the inverse RNA strand exchange assay the RNA concentration was gradually increased at a fixed RPA concentration. We found that at low RNA concentrations, the reaction yield was strongly stimulated by RPA reaching a maximum at 171.5 nM RNA (Figures 6A,B). Further increase in the RNA concentration did not cause an additional increase in the reaction product. In parallel, we determined the stoichiometry of RPA binding to RNA under the same conditions and found that at the concentrations 171.5 nM and below all RNA was sequestered in RPA-RNA complexes and free RNA was only observed at higher RNA concentrations (Figures 6C,D). Thus, the yield of the reaction was increasing with an increase of the RPA-RNA complex concentration but did not show dependence on the presence of free RNA (Figure 6). In contrast, in the absence of RPA, the yield of the reaction promoted by RAD52 was continuously increasing with the increasing of RNA concentration (Figures 6A,B), but not achieving the efficiency observed in the presence of RPA. We concluded that free RNA is not required for the reaction occurring in the presence of RPA, and that RPA acts as a shuttle by delivering RNA to RAD52 for strand exchange with dsDNA.

**Figure 6.**
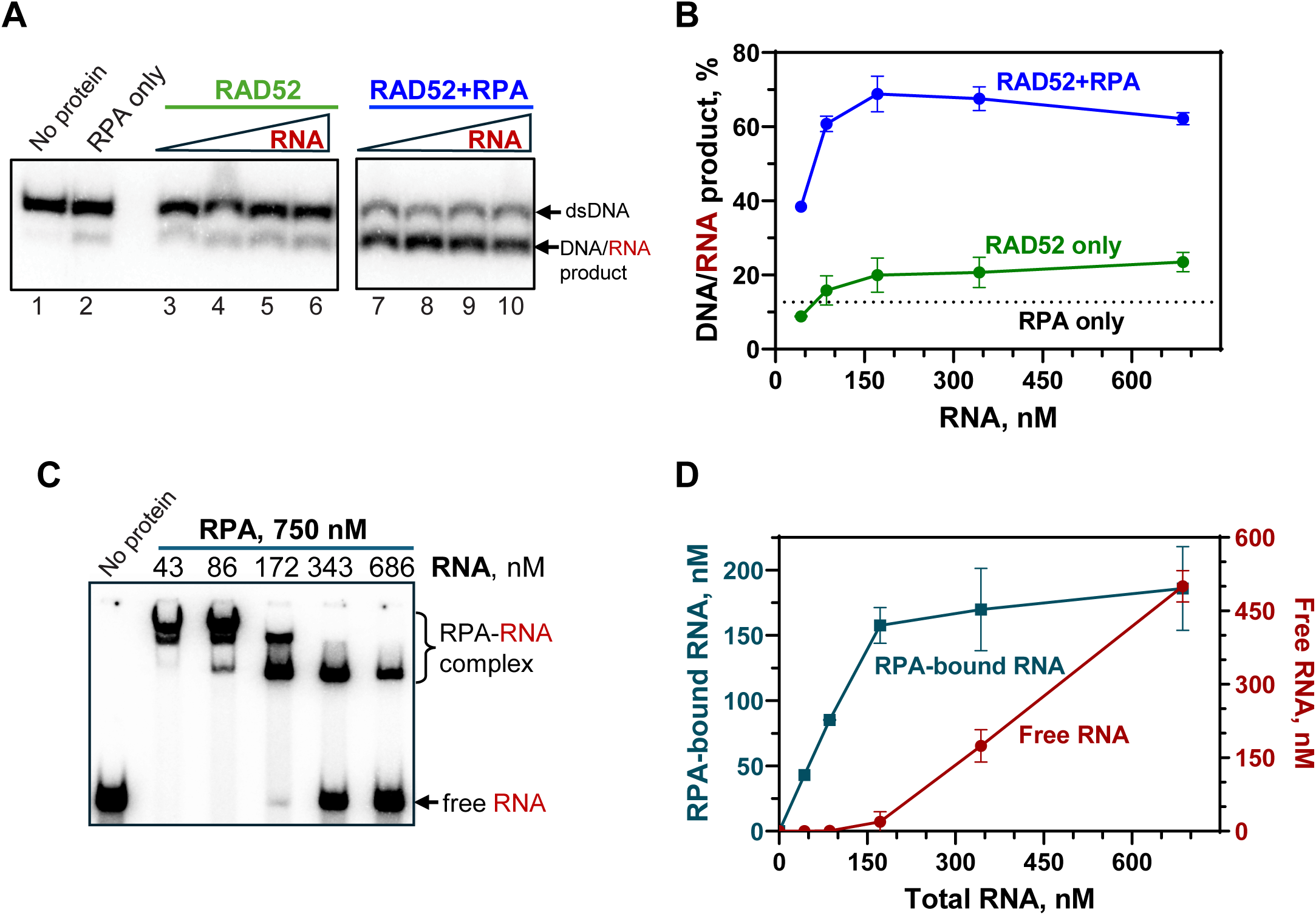
The RPA-RNA complex is an active species in stimulation of RAD52 inverse RNA strand exchange. (**A**) The product formation of inverse RNA strand exchange by RAD52 (1.1 µM) in the absence or presence of RPA (750 nM) in response to increasing RNA (no. 517) concentrations analyzed by electrophoresis in 8% polyacrylamide gels. The RNA concentrations were: 43.2 nM, 85.8 nM, 171.5 nM, 243 nM and 686 nM. **(B)** The data in A presented as a graph. (**C**) A representative polyacrylamide gel image showing the RPA binding to ^32^P-labeled RNA (no. 517). (**D**) The data in (C) plotted as a graph. The experiments were repeated at least 3 times; error bars indicate SD.

## DISCUSSION

In yeast, Rad52 plays a key role in virtually all HR events. In mammals, as evidenced by RAD52 synthetic lethality relationships with the members of the BRCA1/2 protein axis, RAD52 has acquired a role in alternative or back-up HR mechanism(s) (Jalan, Olsen et al. 2019, Rossi, DiDomenico et al. 2021). Particularly, RAD52 appeared to be a central player in RNA-dependent DSB repair (Keskin, Shen et al. 2014, Wei, Nakajima et al. 2015, Mazina, Keskin et al. 2017, Welty, Teng et al. 2018, Yasuhara, Kato et al. 2018). In vitro, RAD52 promotes inverse RNA strand exchange between dsDNA and RNA generating DNA-RNA hybrids. Genetic studies in yeast demonstrated the important role of this activity in RNA-dependent DSB repair (Mazina, Keskin et al. 2017).

We previously found that inverse RNA strand exchange is greatly stimulated by RPA. The RAD52-RPA interaction is evolutionarily conserved and plays a critical role in Rad52 functions in yeast (Plate, Hallwyl et al. 2008). Here, we characterized the role of RAD52-RPA interaction in inverse RNA strand exchange.

Previously, we found that RPA binds to RNA (Mazina, Somarowthu et al. 2020). It was reported that RPA remains accessible for protein-protein interactions when its major DNA binding domains, RPA70A and B, are bound to ssDNA (Brosey, Chagot et al. 2009, Pretto, Tsutakawa et al. 2010, Brosey, Soss et al. 2015). Our current data show that RPA binding to RNA not only preserves but enhances its binding to the RAD52 CTD. RNA cannot act as a bridge in this interaction as the RAD52 CTD does not bind RNA (Figure 1). It is conceivable that conformational changes induced in RPA upon RNA or ssDNA binding cause an increase of the RPA affinity for RAD52 via protein-protein interactions.

Using NMR, we identified two distinct regions of RAD52 CTD involved in RPA binding, the primary (247-280) and secondary (308-335) site. The primary site that overlaps with the previously reported RPA-interacting region in RAD52 (221-280) (Park, Ludwig et al. 1996) and is within the region that NMR observables indicate forms a stable ⍺-helix. The RQK motif in the primary site mediates interactions with the RPA32C subunit for RAD52 and other DNA repair proteins, including UNG2 and XPA (Mer, Bochkarev et al. 2000, Grimme, Honda et al. 2010). Here we showed that the RQK motif is also important for stimulation of RAD52 inverse RNA strand exchange activity by RPA. Further, a previously unknown secondary site in the RAD52 CTD that interacts with RPA70N is also important for the stimulation of inverse RNA strand exchange albeit contributing less than the RQK-containing site (Figure 5B). NMR data indicate that while the second site may have some helical propensity, it remains predominantly disordered under our experimental conditions. It remains possible that the residues in the second site adopt a stable helical conformation upon interaction with RPA70N. Together, this suggests that the largely hydrophobic secondary site could play an important role in the overall stability of the RAD52-RPA complex through a multivalent binding involving binding of short motifs to different regions of the protein partner (Orand, Delaforge et al. 2025). Our studies reveal a more complex mechanism of RAD52-RPA interactions mediated by at least two sites in the CTD that enhance RNA strand exchange activity and speak to potential subtleties in the regulation of this interaction.

Based on our observations, we propose a model of the inverse RNA strand exchange stimulation that involves elaborate cooperation between RAD52 and RPA. We found that the RPA-RNA complex is responsible for the observed stimulation of RAD52 activity, and the formation of this complex serves two important functions (Figure 7).

**Figure 7.**
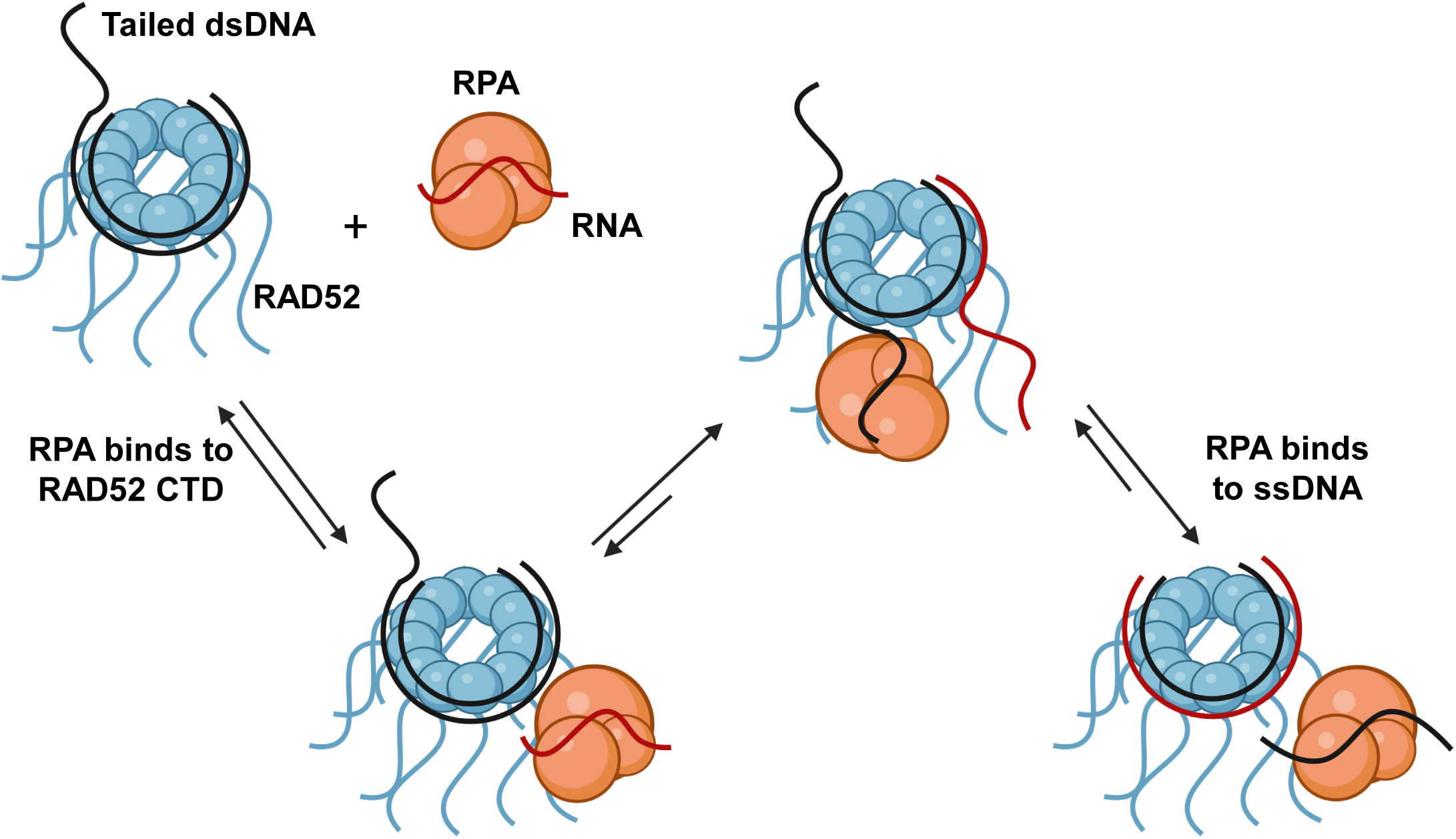
Model for RAD52 inverse RNA strand exchange stimulation by RPA. RPA-RNA (red line) complex interacts with the RAD52-dsDNA (black lines) complex through the two binding sites on the RAD52 CTD. RNA delivered to RAD52 by RPA initiates strand exchange with dsDNA bound to RAD52 producing RNA-DNA hybrid and ssDNA displaced strand intermediates. RPA binding to the displaced ssDNA strand due to its high affinity for ssDNA helping to stabilize the reaction intermediates driving the reaction forward.

First, it strengthens RPA interactions with RAD52 CTD through multiple interaction interfaces and possibly through conformational changes induced in RPA by RNA. Second, binding the RPA-RNA complex to RAD52-dsDNA leads to an increase in the homologous RNA concentration in the proximity of RAD52, which stimulates strand exchange. Inverse RNA strand exchange leads to formation of the displaced ssDNA strand. RPA, due its high affinity for ssDNA, could bind to the displaced strand to further stabilize the reaction intermediate, as it was previously demonstrated for E. coli SSB, the prokaryotic homolog of RPA, in stimulation of DNA strand exchange promoted by RecA (Lavery and Kowalczykowski 1992). Overall, these results provide an example of RNA-induced cooperation between RAD52 and RPA to enhance the inverse RNA strand exchange activity of RAD52 that plays a role in RNA-dependent DNA repair.

## MATERIALS AND METHODS

### DNA and RNA

The DNA and RNA oligonucleotides were purchased from IDT inc. and further purified by electrophoresis in polyacrylamide gels containing 50% urea (Rossi, Mazina et al. 2010). dsDNA substrates were prepared by annealing equimolar (molecule) amounts of indicated complementary oligonucleotides (Rossi, Mazina et al. 2010). When indicated, oligonucleotides were 5’-end labeled with ^32^P using T4 polynucleotide kinase (New England Biolabs) and [γ-^32^P]-ATP. All DNA and RNA concentrations are expressed in moles of molecules, unless indicated otherwise. All experiments with RNA were performed in the presence of 1X Ambion RNA*secure* RNAse inactivation reagent (Mazina, Keskin et al. 2017).

### Purification of RAD52 CTD wild type, QEM, and ANQ mutants

The pGEX-6P-2 plasmid with GST-RAD52-CTD (217-418) wild type or mutants (QEM or ANQ) was transformed into *E. coli* BL21 (DE3) cells (New England Biolabs, MA). Cells were grown in LB with 100 µg/mL Ampicillin at 37°C until an OD_600_ of 0.6 was reached. Then, protein expression was induced by 0.4 mM IPTG and cells were incubated for 4 h at 37°C. All purification steps were carried out at 4°C. Harvested cells were resuspended in Lysis Buffer (40 mM KH_2_PO_4_, pH 7.5, 500 mM KCl, 5 mM EDTA, 10% glycerol, 14.3 mM β-mercaptoethanol,1 mM PMSF) supplemented with EDTA-free protease inhibitor cocktail (Roche Applied Science) and lysed by passing cell suspension through Emulsiflex-C3 (Avestin) three times and centrifuged for 1 h at 40,000 rpm (50.2 Ti rotor, Beckman) at 4°C. Supernatant was loaded on to a 10 mL Glutathione Sepharose 4 FF (Cytiva) column, washed with 5 column volumes of PBS buffer (8.1 mM Na_2_HPO_4_, 1.47 mM KH_2_PO_4_, 138 mM NaCl, 2.7 mM KCl pH 7.4) supplemented with 500 mM KCl and 14.3 mM β-mercaptoethanol, and 5 column volumes of Lysis Buffer before being eluted with GST Elution Buffer (50 mM Tris-HCl, pH 8.0, 500 mM KCl, 1 mM EDTA, 20 mM glutathione, 10 % glycerol, 14.3 mM β - mercaptoethanol). Fractions containing RAD52 CTD were pooled and concentrated using an Amicon concentrator (30 kDa MWCO) before being loaded onto a 60 mL Superdex 200 column equilibrated with K_20_Cl_300_ buffer (20 mM KH_2_PO_4_, pH 7.5, 300 mM KCl, 10% glycerol, 14.3 mM β-mercaptoethanol). For NMR experiments, the GST tag was cleaved by incubation of pooled fractions containing RAD52 CTD with 1 µg of PreScission Protease (Cytiva) per 100 µg of total protein overnight on ice. Then, GST-RAD52 or GST-cleaved RAD52 sample was diluted to 100 mM KCl using dilution buffer (50 mM KH_2_PO_4_, pH 7.5, 10% glycerol, and 14.3 mM β-mercaptoethanol) and loaded onto a 5 mL Heparin column, followed by a 5 column volume wash and fractionated using a 30 column volume gradient of 30-400 mM KCl in buffer K_20_Cl_30_ (20 mM KH_2_PO_4_, pH 7.5, 10% glycerol, 14.3 mM β-mercaptoethanol). Fractions were tested for nuclease contamination using ^32^P-labeled ssDNA as a substrate. Nuclease-free fractions were combined and concentrated using an Amicon concentrator (10 kDa MWCO), dialyzed against NMR buffer (20 mM NaH_2_PO_4_, pH 6.8, 100 mM NaCl, 1 mM DTT) using Spectra Pore Float-a-lyzer G2 dialysis device overnight at 4°C, and concentrated again using an Amicon concentrator (10 kDa MWCO). All purified proteins were aliquoted and stored at – 80°C.

### 15N or ^13^C-RAD52 CTD purification for NMR studies

The *E. coli* cells carrying pGEX-6P-2 plasmid with GST-RAD52-CTD (217-418) were grown at 37°C in M9 medium that contained ^15^NH_4_Cl (≥98 atom % ^15^N; Sigma Isotec #299251) 1 g/L, and ^13^C_6_-D-glucose (Sigma Isotec #389374) 3 g/L (for 3D assignment experiments only), as the sole nitrogen and carbon source, respectively. The medium was supplemented with 10% (v/v) MEM Vitamin Solution (ThermoFisher, #MT25020CI), 0.02% (w/v) yeast extract, and 100 µg/mL Ampicillin. Protein expression was induced at OD_600_ ∼ 0.6 by the addition of 0.4 mM IPTG, growth continued for 5 h at 37°C. The ^15^N or ^15^N,^13^C-labeled RAD52 CTD was purified using the same scheme as for unlabeled RAD52 CTD with the addition step of GST tag removal using 1 µg of PreScission Protease (Cytiva) per 100 µg of total protein incubated overnight on ice before loading onto the heparin column.

### Purification of full-length wild type and mutant RAD52 proteins

The pGEX-6P-2 plasmid encoding GST-RAD52 wild type or mutants (GST-RAD52-Δ251-418, GST-RAD52-QEM, GST-RAD52-ANQ, GST-RAD52-QEM+ANQ) was transformed into *E. coli* BL21 (DE3) cells (New England Biolabs). Cells were grown in LB at 37°C until an OD_600_ of 0.6 was reached and then protein expression was induced by 0.4 mM IPTG followed by incubation for 16 h at 16°C. All purification steps were carried out at 4°C. Harvested cells were resuspended in Lysis Buffer (50 mM Tris-HCl, pH 7.5, 200 mM KCl, 1 mM EDTA, 10% glycerol, 14.3 mM β-mercaptoethanol,1 mM PMSF) supplemented with EDTA-free protease inhibitor cocktail (Roche Applied Science) and lysed by passing cell suspension through the EmulsiFlex-C3 (Avestin) three times and centrifuged for 1 h at 30,000 rpm (50.2 Ti rotor, Beckman) at 4°C. Supernatant was loaded on to a 20 mL Q Sepharose column equilibrated with lysis buffer. The flow through was subjected to ammonium sulfate precipitation at 40% saturation and centrifuged at 12,000 rpm (JA-25.50) for 15 min at 4°C. The pellet was resuspended in 80 mL of lysis buffer and loaded onto a Glutathione Sepharose 4 FF (Cytiva) column. The column was washed with 5 column volumes of PBS buffer (1X Phosphate buffered saline, 500 mM KCl, 14.3 mM β-mercaptoethanol) and 5 column volumes of T_50_Cl_500_ (50 mM Tris-HCl, pH 7.5, 500 mM KCl, 1 mM EDTA, 10% glycerol, 14.3 mM β-mercaptoethanol) before being eluted with GST Elution Buffer (50 mM Tris-HCl, pH 8.0, 500 mM KCl, 1 mM EDTA, 20 mM glutathione, 10 % glycerol, 14.3 mM β - mercaptoethanol). Fractions containing RAD52 were pooled and incubated with PreScission Protease (Cytiva) (1 µg per 100 µg of total protein) overnight on ice.

Following cleavage of the GST tag, RAD52 was loaded onto a 5 mL Heparin column equilibrated with K_50_Cl_100_ buffer (50 mM KH_2_PO_4_, pH 7.5, 100 mM KCl, 10% glycerol, and 14.3 mM β-mercaptoethanol) followed by a 5 column volume wash with K_50_Cl_100_ buffer and developed with a 20 column volume gradient of 100-1200 mM KCl in buffer K_50_Cl_100_ (50 mM KH_2_PO_4_, pH 7.5, 100 mM KCl, 10% glycerol, 14.3 mM β-mercaptoethanol). Fractions containing RAD52 were combined and diluted to 100 mM KCl with buffer K_50_ (50 mM KH_2_PO_4_, pH 7.5, 10% glycerol, 14.3 mM β-mercaptoethanol) and loaded onto a 1 mL Mono S 5/50 GL column equilibrated with K_50_Cl_100._ The sample was washed with 5 column volumes of K_50_Cl_100_ before fractionating with a 20 column-volume gradient of 100-600 mM KCl in buffer (50 mM KH_2_PO_4_, pH 7.5, 100 mM KCl, 10% glycerol, 14.3 mM β-mercaptoethanol). Fractions containing RAD52 were tested for nuclease contamination using ^32^P-labeled ssDNA as a substrate. Nuclease-free fractions were combined and dialyzed using Spectra Pore Float-a-lyzer G2 dialysis device against storage buffer (50 mM Tris-HCl, pH 7.5, 200 mM NaCl, 15% glycerol, 1 mM DTT) overnight at 4°C.

### Purification of human RPA

The p11d-tRPA plasmid was transformed into E. coli BL21 (DE3) cells (New England Biolabs). Cells were grown in TB (10 g Bacto-Tryptone, 5 g NaCl per 1 L) at 37°C until an OD_600_ of 0.6 was reached and protein expression was induced by 0.4 mM IPTG for 2 h at 37°C. Cells from 12 L of culture were resuspended in HEPES:NaOH Lysis Buffer (30 mM HEPES:NaOH, pH 7.8, 5 mM EDTA, 10% sucrose, 0.01% NP-40, 20 mM β-mercaptoethanol) supplemented with 1 mM PMSF and EDTA-free protease inhibitor cocktail (Roche Applied Science). Cells were lysed by passing cell suspension through the EmulsiFlex-C3 (Avestin) three times and centrifuged for 1 h at 40,000 rpm (50.2 Ti rotor, Beckman) at 4°C. For RPA purification we followed the procedure described in (Henricksen, Umbricht et al. 1994). Supernatant was applied to a 70 mL Affi-Gel blue column equilibrated with HI buffer (30 mM HEPES:NaOH, pH 7.8, 1 mM EDTA, 0.25% inositol, 0.01% NP-40, 10 mM β-mercaptoethanol) containing 50 mM KCl. The column was washed sequentially with HI buffer containing 50 mM KCl, 0.8 M KCl, and 0.5 M NaSCN. RPA was eluted with an 8-column volume gradient of 500 mM-1.5 M NaSCN in buffer HI +NaSCN_500_ (30 mM HEPES:NaOH, pH 7.8, 500 mM NaSCN, 0.25 mM EDTA, 0.25% inositol, 0.01% NP-40, 10 mM β-mercaptoethanol). Peak fractions containing RPA were pooled and applied to a hydroxylapatite column (15 mL) equilibrated with HI buffer and then eluted with 4 column volumes of HI buffer containing 80 mM potassium phosphate. The peak fractions were collected and diluted with an equal volume of T_25_ buffer (25 mM Tris-HCl, pH 7.5, 0.1 mM EDTA, 10% glycerol, 10 mM β-mercaptoethanol) and filtered through at 0.2 µM PVDF filter before being applied to a 1 mL Mono-Q column. The sample was washed with 3 mL of HI buffer containing 100 mM KCl and developed with a 20 column-volume gradient 100-600 mM KCl in T_25_ buffer.

Fractions were tested for nuclease activity against ^32^P-labeled ssDNA. Nuclease-free fractions were combined and concentrated using an Amicon concentrator (10 kDa MWCO), dialyzed against NMR buffer (20 mM NaH_2_PO_4_, pH 6.8, 100 mM NaCl, 1 mM DTT) using Spectra Pore Float-a-lyzer G2 dialysis device overnight at 4°C, and concentrated again using an Amicon concentrator (10 kDa MWCO). For biochemical experiments, RPA was dialyzed against storage buffer (25 mM Tris-HCL, pH 7.5, 300 mM KCl, 0.1 mM EDTA, 1 mM DTT, 30% glycerol).

Human RPA32C (202-270) and RPA70N (1-120) constructs were cloned in pET15b (Novagen) and overexpressed in the E. coli strain BL21(DE3) pLysS and purified as described in (Mer, Bochkarev et al. 2000).

### Nuclear Magnetic Resonance Spectroscopy

All experiments were recorded on a Bruker Avance NEO spectrometer operating at a proton Larmor frequency of 700.05 MHz, at a temperature of 298 K. Data were processed with Topspin 4.4.0 (Bruker) or NMRPipe (Delaglio, Grzesiek et al. 1995) and analyzed with CCPNMR Analysis 2.5.2 software (Skinner, Fogh et al. 2016). RAD52 CTD (217-418) ^1^H, ^13^Cα, ^13^Cβ, ^13^C’ and ^15^N backbone resonances were assigned through the analysis of a set of 2D and 3D experiments, including ^1^H,^15^N-HSQC, HNCACB, CBCA(CO)NH, HNCO, HN(CA)CO and HNCC(CO)NH, recorded on a ^13^C, ^15^N-labeled RAD52 CTD (217-418) sample at a concentration of 130 μM, in NMR buffer (20 mM sodium phosphate pH 6.8, 100 mM sodium chloride, 1 mM DTT). The ^1^H,^15^N-HSQC was recorded with 128*x 1024* complex points in the indirect (^15^N) and direct (^1^H) dimensions, corresponding to acquisition times of 75.2 and 112.6 ms, respectively. Acquisition parameters for the HNCO and HN(CA)CO consisted of 32* x 64* x 1024* complex points in the indirect (F1, ^13^C), (F2, ^15^N) and direct (F3 ^1^H) dimensions, corresponding to acquisition times of 15.2, 37.6, 112.6 ms, respectively; acquisition parameters for the HNCACB, CBCA(CO)NH and HNCC(CO)NH consisted of 112* x 64* x 1024* complex points in the indirect (F1, ^13^C), (F2, ^15^N) and direct (F3 ^1^H) dimensions, corresponding to acquisition times of 9.6, 37.6, 112.6 ms, respectively. All 3D experiments were recorded in non-uniform sampling (NUS) mode (Delaglio, Walker et al. 2017) with a sampling density of 20%, and the spectra were reconstructed using the SMILE algorithm implemented in NMRPipe (Ying, Delaglio et al. 2017). The proportion of secondary structure was estimated from the chemical shifts of the backbone atoms ^1^H_N_, ^15^N, ^13^C⍺, ^13^Cβ, and ^13^C’ using the SSP algorithm (Marsh, Singh et al. 2006). The ^15^N longitudinal *R*_1_ and transverse *R*_2_ relaxation rates were calculated from *T*_1_ and *T*_1ρ_ experiments (Lakomek, Ying et al. 2012), recorded on 50 µM sample of ^15^N RAD52 CTD (217-418) in NMR buffer using 64* x 1024* complex data points in the indirect (^15^N) and direct (^1^H) dimensions corresponding to acquisition times of 37.6 and 112.6 ms, respectively. The ^15^N *T*_1_ experiment consisted of eight interleaved spectra with the following relaxation delays: 40, 80, 200, 280, 300, 400, 600, and 800 ms. The *T*_1ρ_ experiment was recorded using a B_1_ field of 1400 Hz and eight interleaved spectra with the following relaxation delays: 1, 21, 31, 41, 61, 81, 121 and 161 ms. The ^15^N *R*_2_ rates were calculated using the following equation (Massi, Johnson et al. 2004).

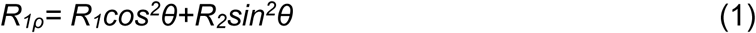

with θ = arctan(ω_1_/Ω), where ω_1_ is the B_1_ field strength (here 1400 Hz) and Ω is the offset from the spinlock carrier frequency. ^1^H-^15^N heteronuclear NOE experiments were recorded on the same sample and consisted of two interleaved experiments, with and without proton saturation, using a recycle delay of 4 s. Spectra were acquired with 64* x 1024* complex data points in the indirect (^15^N) and direct (^1^H) dimensions corresponding to acquisition times of 37.6 and 112.6 ms, respectively.

For all titration experiments RAD52 CTD (217-418) was uniformly ^15^N labeled and the RPA heterotrimer, RPA32C, and RPA70N, were the unlabeled (e.g. ^14^N) titrant. ^1^H,^15^N-HSQC spectra were recorded at 298 K, with 64* x 1024* complex data points in the indirect (^15^N) and direct (^1^H) dimensions, corresponding to acquisition times of 37.6 and 112.6 ms, respectively. For the RPA heterotrimer, a 200 µL mixture of 35 µM ^15^N RAD52-CTD and 35 µM ^14^N RPA FL was prepared, 32.4 and 70 µL aliquots of 216 µM ^15^N RAD52-CTD were added for RAD52:RPA ratios of 1:1, 2:1, and 4:1, respectively.

Changes in peak intensities were calculated using a 50 µM sample of ^15^N RAD52-CTD suspended in identical buffer as a reference.

For RPA32C, aliquots of 1 mM RPA32C were titrated into a 50 µM ^15^N RAD52-CTD sample in 160 µL of NMR buffer in a 3 mm NMR tube (Wilmad, NJ) resulting in final RAD52:RPA32C ratios of 4:1, 2:1, and 1:1. For RPA70N, total of 50 µL (in aliquots) of 1 mM RPA70N were titrated into a sample of 50 µM ^15^N RAD52-CTD in 500 µL of NMR buffer in a 5 mm NMR tube (Wilmad, NJ) for final RAD52:RPA70N ratios of 4:1, 2:1, 1:1, and 1:2. Spectra were apodized with a sine bell function and zero-filled to twice the number of acquired points for data analysis by CCPNMR 2.5.2 software. Line broadening analysis was performed by calculating the change in peak intensity (*I/I*₀) between RAD52-CTD and the highest RPA:RAD52 ratio recorded (Waudby, Ramos et al. 2016). Chemical shift perturbations (CSP) were calculated by weighting the ^1^H and ^15^N chemical shifts according to their gyromagnetic ratios using the following equation (Williamson 2013).

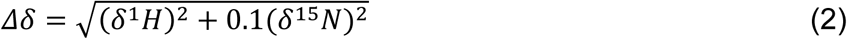

CSPs were considered significant when they were higher than the sum of average and standard deviation of Δδ_max_ for all residues.

### *In vitro* pull-downs

Glutathione Sepharose 4 Fast Flow resin (Cytiva) was pre-equilibrated in binding buffer (20 mM KH_2_PO_4_, pH 7.4, 100 mM KCl, 10% glycerol, 0.01% NP-40). A reaction mixture containing 6 µM RPA and 6 µM ssDNA (no. 211, 48-mer) or 6 µM RNA (no. 517, 63-mer) was preincubated for 15 min at 37°C before adding 6 µM of GST-RAD52-CTD, GST-RAD52-CTD-QEM, or GST-RAD52-CTD-ANQ in a final volume of 30 µL of binding buffer and incubated on ice. After 1 h, the reaction mixture was added to the resin (30 µL of slurry) and incubated on ice for 1 h. Samples were centrifuged for 30 sec on a tabletop centrifuge (USA Scientific; 2500 x g) and the supernatant was collected. Resin was washed three times with 150 µL of binding buffer. The bound fraction was eluted with 28 µL of elution buffer (50 mM Tris-HCl, pH 8.0, 20 mM glutathione, 100 mM NaCl).

Fractions were analyzed by electrophoresis in 12% SDS-polyacrylamide gels (SDS-PAGE). Gel was stained with Coomassie blue.

### RAD52 or RAD52 CTD binding to ssDNA or RNA using electrophoretic mobility shift assay (EMSA)

RAD52 or RAD52-CTD (0.5, 1, 1.5, or 2 µM) were incubated with ssDNA (no. 1, 63-mer) or RNA (no. 517, 63-mer) (63 nM) in buffer containing 20 mM HEPES-NaOH, pH 7.5, 2 mM β-mercaptoethanol, and 0.01 mg/mL bovine serum albumin. The reaction mixture was incubated for 10 min at 37°C, and then 5 µL of 70% glycerol, 1% bromophenol blue was added. Samples were analyzed by electrophoresis in 8% polyacrylamide gels (acrylamide: N, N’-methylenebisacrylamide, 29:1) in 1X TBE buffer (89 nM Tris, 89 nM boric acid and 1 mM EDTA, pH 8.0) at 13 V/cm for 1 h at room temperature. The gels were dried on Amersham Hybond-N + membrane (Cytiva) and analyzed using a Typhoon RGB biomolecular imager with the Image QuantTL software (Cytiva).

### RPA binding to RNA using EMSA

Nucleoprotein complexes were assembled by incubating RPA (750 nM) with ^32^P-labeled RNA (no. 517) in indicated concentrations in a buffer A (25 mM Tris-Acetate, pH 7.5, 100 µg/µL BSA, 2 mM magnesium acetate, and 2 mM DTT) for 5 min at 37°C. The samples (10 µL) were mixed with 1.5 µl of 50% glycerol and loaded onto a 6% polyacrylamide (acrylamide: N, N’-methylenebisacrylamide, 29:1). 1.5 uL of 70% glycerol and 0.01% Bromphenol blue was added only in the sample containing ^32^P-labeled RNA probe without RPA. Samples were analyzed by electrophoresis using 0.25 X TBE buffer (89 nM Tris, 89 nM boric acid and 1 mM EDTA, pH 8.0) at 13V/cm for 1 h at room temperature. The gels were dried on Amersham Hybond-N + membrane (Cytiva) and analyzed using a Typhoon RGB biomolecular imager with the Image QuantTL software (Cytiva).

### Inverse RNA strand exchange assays

Nucleoprotein complexes were assembled by incubating RAD52 (1.1 µM) with ^32^P-labeled 3’-tailed dsDNA (no.1/ no. 117; 68.6 nM) in a buffer A containing 25 mM Tris-Acetate, pH 7.5, 100 µg/µL BSA, 2 mM magnesium acetate, and 2 mM DTT for 15 min at 37°C. The reactions were initiated by the addition of RNA (no. 517; 686 nM). Aliquots (10 µL) were withdrawn at the indicated time points and DNA or RNA samples were deproteinized by incubation in 1% SDS, 1.6 mg/mL proteinase K, 6% glycerol and 0.01% bromophenol blue for 15 min at 37°C. Samples were analyzed by electrophoresis in 8% polyacrylamide gels (acrylamide: N, N’-methylenebisacrylamide, 29:1) in 1X TBE buffer (89 nM Tris, 89 nM boric acid and 1 mM EDTA, pH 8.0) at 13V/cm for 1 h at room temperature. The gels were dried on Amersham Hybond-N + membrane (GE Healthcare) and analyzed using a Typhoon RGB biomolecular imager with the Image QuantTL software (GE Healthcare).

When human RPA (1.5 µM) was used, it was preincubated with RNA (no. 517; 1.37 µM) in buffer A for 15 min at 37°C. A separate reaction mixture containing RAD52 (2.2 µM) and labeled 3’ tailed DNA (137 nM) in buffer A was incubated for 15 min at 37°C. Inverse strand exchange reaction was initiated by addition of mixtures in a 1:1 ratio containing RPA and RNA to the RAD52 nucleoprotein complexes. The final concentration of RPA was 750 nM.

The conditions for the mutants were the same, except the final protein concentrations for each mutant were as follows, NTD (1.4 µM), Δ251-418 (2 µM), QEM (2 µM), ANQQAA (1.7 µM), QEM-ANQQAA (1.5 µM). The conditions for the RPA domain constructs were the same, except the final protein concentrations were RPA70N (0, 0.4, 0.75, 1.2, 1.5, 2 µM), RPA32C (0, 0.4, 0.75, 1.2, 1.5, 2 µM), and RPA70N:RPA32C (1:1) (0, 0.75, 1.5, 2, 3.5, 5.5 µM, of each domain). The optimal concentrations of RAD52 and RAD52 mutant proteins were determined in preliminary experiments.

## Acknowledgments

We are thankful to Dr. Courtney Johnson for her comments and suggestions.

## Data Availability

“Assignment of ^1^H, ^13^C, ^15^N backbone chemical shifts of RAD52 217-418” were deposited in the BMRB under accession number 53286.

## Author contributions

Conceptualization: SFD, MJR, DSL, AVM Investigation: SFD, MJR, HHD, AB, SSS.

Generated requisite materials including plasmids, DNA, RNA, and purified proteins: SFD, MJR, HHD, AB, WJC

Structural Analysis: HHD, AB, DSL

Analyzed data and wrote the manuscript: SFD, MJR, HHD, WJC, DSL, AVM

## REFERENCES

Balboni, B., R. Marotta, F. Rinaldi, G. Milordini, G. Varignani, S. Girotto and A. Cavalli (2024). “An integrative structural study of the human full-length RAD52 at 2.2 A resolution.” Commun Biol 7(1): 956.

Balboni, B., F. Rinaldi, V. Previtali, A. Ciamarone, S. Girotto and A. Cavalli (2023). “Novel Insights into RAD52’s Structure, Function, and Druggability for Synthetic Lethality and Innovative Anticancer Therapies.” Cancers (Basel) 15(6).

Baudin, A., H. H. Dinh, X. Xu and D. S. Libich (2025). “The (1)H, (15)N and (13)C backbone resonance assignments of the N-terminal (1-149) domain of Serpine mRNA Binding Protein 1 (SERBP1).” Biomol NMR Assign 19(1): 101–107.

Bochkareva, E., L. Kaustov, A. Ayed, G. S. Yi, Y. Lu, A. Pineda-Lucena, J. C. Liao, A. L. Okorokov, J. Milner, C. H. Arrowsmith and A. Bochkarev (2005). “Single-stranded DNA mimicry in the p53 transactivation domain interaction with replication protein A.” Proc Natl Acad Sci U S A 102(43): 15412–15417.

Brosey, C. A., M. E. Chagot, M. Ehrhardt, D. I. Pretto, B. E. Weiner and W. J. Chazin (2009). “NMR analysis of the architecture and functional remodeling of a modular multidomain protein, RPA.” J Am Chem Soc 131(18): 6346–6347.

Brosey, C. A., S. E. Soss, S. Brooks, C. Yan, I. Ivanov, K. Dorai and W. J. Chazin (2015). “Functional dynamics in replication protein A DNA binding and protein recruitment domains.” Structure 23(6): 1028–1038.

Chandramouly, G., J. Zhao, S. McDevitt, T. Rusanov, T. Hoang, N. Borisonnik, T. Treddinick, F. W. Lopezcolorado, T. Kent, L. A. Siddique, J. Mallon, J. Huhn, Z. Shoda, E. Kashkina, A. Brambati, J. M. Stark, X. S. Chen and R. T. Pomerantz (2021). “Poltheta reverse transcribes RNA and promotes RNA-templated DNA repair.” Sci Adv 7(24).

Delaglio, F., S. Grzesiek, G. W. Vuister, G. Zhu, J. Pfeifer and A. Bax (1995). “NMRPipe: a multidimensional spectral processing system based on UNIX pipes.” J Biomol NMR 6(3): 277–293.

Delaglio, F., G. S. Walker, K. A. Farley, R. Sharma, J. C. Hoch, L. W. Arbogast, R. G. Brinson and J. P. Marino (2017). “Non-Uniform Sampling for All: More NMR Spectral Quality, Less Measurement Time.” Am Pharm Rev 20(4).

Fan, J. and N. P. Pavletich (2012). “Structure and conformational change of a replication protein A heterotrimer bound to ssDNA.” Genes Dev 26(20): 2337–2347.

Feng, Z., S. P. Scott, W. Bussen, G. G. Sharma, G. Guo, T. K. Pandita and S. N. Powell (2011). “Rad52 inactivation is synthetically lethal with BRCA2 deficiency.” Proc Natl Acad Sci U S A 108(2): 686–691.

Game, J. C. and R. K. Mortimer (1974). “A genetic study of X-ray sensitive mutants in yeast.” Mutation Research 24: 281.

Grimme, J. M., M. Honda, R. Wright, Y. Okuno, E. Rothenberg, A. V. Mazin, T. Ha and M. Spies (2010). “Human Rad52 binds and wraps single-stranded DNA and mediates annealing via two hRad52-ssDNA complexes.” Nucleic Acids Res 38(9): 2917–2930.

Hanahan, D. and R. A. Weinberg (2011). “Hallmarks of cancer: the next generation.” Cell 144(5): 646–674.

Hanamshet, K., O. M. Mazina and A. V. Mazin (2016). “Reappearance from Obscurity: Mammalian Rad52 in Homologous Recombination.” Genes (Basel) 7(9).

Henricksen, L. A., C. B. Umbricht and M. S. Wold (1994). “Recombinant replication protein A: expression, complex formation, and functional characterization.” J Biol Chem 269(15): 11121–11132.

Honda, M., M. Razzaghi, P. Gaur, E. Malacaria, G. Marozzi, L. Di Biagi, F. A. Aiello, E. A. Paintsil, A. J. Stanfield, B. J. Deppe, L. Gakhar, N. J. Schnicker, M. A. Spies, P. Pichierri and M. Spies (2025). “The RAD52 double-ring remodels replication forks restricting fork reversal.” Nature 641(8062): 512–519.

Jackson, D., K. Dhar, J. K. Wahl, M. S. Wold and G. E. Borgstahl (2002). “Analysis of the human replication protein A:Rad52 complex: evidence for crosstalk between RPA32, RPA70, Rad52 and DNA.” J Mol Biol 321(1): 133–148.

Jalan, M., A. Brambati, H. Shah, N. McDermott, J. Patel, Y. Zhu, A. Doymaz, J. Wu, K. S. Anderson, A. Gazzo, F. Pareja, T. N. Yamaguchi, T. Vougiouklakis, S. Ahmed-Seghir, P. Steinberg, A. Neiman-Golden, B. Azeroglu, J. Gomez-Aguilar, E. M., da Silva, S. Hussain, D. Higginson, P. C. Boutros, N. Riaz, J. S. Reis-Filho, S. N. Powell and A. Sfeir (2025). “RNA transcripts serve as a template for double-strand break repair in human cells.” Nat Commun 16(1): 4349.

Jalan, M., K. S. Olsen and S. N. Powell (2019). “Emerging Roles of RAD52 in Genome Maintenance.” Cancers (Basel) 11(7).

Johnson, C. N., X. Xu, S. P. Holloway and D. S. Libich (2022). “The (1)H, (15)N and (13)C resonance assignments of the low-complexity domain from the oncogenic fusion protein EWS-FLI1.” Biomol NMR Assign 16(1): 67–73.

Kagawa, W., A. Kagawa, K. Saito, S. Ikawa, T. Shibata, H. Kurumizaka and S. Yokoyama (2008). “Identification of a second DNA binding site in the human Rad52 protein.” J Biol Chem 283(35): 24264–24273.

Kagawa, W., H. Kurumizaka, S. Ikawa, S. Yokoyama and T. Shibata (2001). “Homologous pairing promoted by the human Rad52 protein.” J Biol Chem 276(37): 35201–35208.

Kagawa, W., H. Kurumizaka, R. Ishitani, S. Fukai, O. Nureki, T. Shibata and S. Yokoyama (2002). “Crystal structure of the homologous-pairing domain from the human Rad52 recombinase in the undecameric form.” Mol Cell 10(2): 359–371.

Keskin, H., Y. Shen, F. Huang, M. Patel, T. Yang, K. Ashley, A. V. Mazin and F. Storici (2014). “Transcript-RNA-templated DNA recombination and repair.” Nature 515(7527): 436–439.

Kinoshita, C., Y. Takizawa, M. Saotome, S. Ogino, H. Kurumizaka and W. Kagawa (2023). “The cryo-EM structure of full-length RAD52 protein contains an undecameric ring.” FEBS Open Bio 13(3): 408–418.

Koike, M., Y. Yutoku and A. Koike (2013). “The C-terminal region of Rad52 is essential for Rad52 nuclear and nucleolar localization, and accumulation at DNA damage sites immediately after irradiation.” Biochem Biophys Res Commun 435(2): 260–266.

Kowalczykowski, S. C. (2015). “An Overview of the Molecular Mechanisms of Recombinational DNA Repair.” Cold Spring Harb Perspect Biol 7(11).

Krogh, B. O. and L. S. Symington (2004). “Recombination proteins in yeast.” Annu Rev Genet 38: 233–271.

Lakomek, N. A., J. Ying and A. Bax (2012). “Measurement of (1)(5)N relaxation rates in perdeuterated proteins by TROSY-based methods.” J Biomol NMR 53(3): 209–221.

Lavery, P. E. and S. C. Kowalczykowski (1992). “A postsynaptic role for single-stranded DNA-binding protein in recA protein-promoted DNA strand exchange.” J Biol Chem 267(13): 9315–9320.

Liang, C. C., L. A. Greenhough, L. Masino, S. Maslen, I. Bajrami, M. Tuppi, M. Skehel, I. A. Taylor and S. C. West (2024). “Mechanism of single-stranded DNA annealing by RAD52-RPA complex.” Nature 629(8012): 697–703.

Libich, D. S., N. L. Fawzi, J. Ying and G. M. Clore (2013). “Probing the transient dark state of substrate binding to GroEL by relaxation-based solution NMR.” Proc Natl Acad Sci U S A 110(28): 11361–11366.

Lloyd, J. A., D. A. McGrew and K. L. Knight (2005). “Identification of residues important for DNA binding in the full-length human Rad52 protein.” J Mol Biol 345(2): 239–249.

Ma, C. J., Y. Kwon, P. Sung and E. C. Greene (2017). “Human RAD52 interactions with replication protein A and the RAD51 presynaptic complex.” J Biol Chem 292(28): 11702–11713.

Marsh, J. A., V. K. Singh, Z. Jia and J. D. Forman-Kay (2006). “Sensitivity of secondary structure propensities to sequence differences between alpha-and gamma-synuclein: implications for fibrillation.” Protein Sci 15(12): 2795–2804.

Massi, F., E. Johnson, C. Wang, M. Rance and A. G. Palmer, 3rd (2004). “NMR R1 rho rotating-frame relaxation with weak radio frequency fields.” J Am Chem Soc 126(7): 2247–2256.

Mazina, O. M., H. Keskin, K. Hanamshet, F. Storici and A. V. Mazin (2017). “Rad52 Inverse Strand Exchange Drives RNA-Templated DNA Double-Strand Break Repair.” Mol Cell 67(1): 19–29 e13.

Mazina, O. M., S. Somarowthu, L. Y. Kadyrova, A. G. Baranovskiy, T. H. Tahirov, F. A. Kadyrov and A. V. Mazin (2020). “Replication protein A binds RNA and promotes R-loop formation.” J Biol Chem.

Meers, C., H. Keskin, G. Banyai, O. Mazina, T. Yang, A. L. Gombolay, K. Mukherjee, E. I. Kaparos, G. Newnam, A. Mazin and F. Storici (2020). “Genetic Characterization of Three Distinct Mechanisms Supporting RNA-Driven DNA Repair and Modification Reveals Major Role of DNA Polymerase zeta.” Mol Cell 79(6): 1037–1050 e1035.

Mer, G., A. Bochkarev, R. Gupta, E. Bochkareva, L. Frappier, C. J. Ingles, A. M. Edwards and W. J. Chazin (2000). “Structural basis for the recognition of DNA repair proteins UNG2, XPA, and RAD52 by replication factor RPA.” Cell 103(3): 449–456.

Mortensen, U. H., C. Bendixen, I. Sunjevaric and R. Rothstein (1996). “DNA strand annealing is promoted by the yeast Rad52 protein.” Proc Natl Acad Sci U S A 93(20): 10729–10734.

Mortensen, U. H., M. Lisby and R. Rothstein (2009). “Rad52.” Curr Biol 19(16): R676– 677.

Orand, T., E. Delaforge, A. Lee, J. Kragelj, M. Tengo, L. Tengo, M. Blackledge, E. Boeri Erba, R. J. Davis, A. Palencia and M. R. Jensen (2025). “Bipartite binding of the intrinsically disordered scaffold protein JIP1 to the kinase JNK1.” Proc Natl Acad Sci U S A 122(9): e2419915122.

Park, M. S., D. L. Ludwig, E. Stigger and S. H. Lee (1996). “Physical interaction between human RAD52 and RPA is required for homologous recombination in mammalian cells.” J Biol Chem 271(31): 18996–19000.

Plate, I., S. C. Hallwyl, I. Shi, L. Krejci, C. Muller, L. Albertsen, P. Sung and U. H. Mortensen (2008). “Interaction with RPA is necessary for Rad52 repair center formation and for its mediator activity.” J Biol Chem 283(43): 29077–29085.

Prakash, A. and G. E. Borgstahl (2012). “The structure and function of replication protein A in DNA replication.” Subcell Biochem 62: 171–196.

Pretto, D. I., S. Tsutakawa, C. A. Brosey, A. Castillo, M. E. Chagot, J. A. Smith, J. A. Tainer and W. J. Chazin (2010). “Structural dynamics and single-stranded DNA binding activity of the three N-terminal domains of the large subunit of replication protein A from small angle X-ray scattering.” Biochemistry 49(13): 2880–2889.

Rijkers, T., J. Van Den Ouweland, B. Morolli, A. G. Rolink, W. M. Baarends, P. P. Van Sloun, P. H. Lohman and A. Pastink (1998). “Targeted inactivation of mouse RAD52 reduces homologous recombination but not resistance to ionizing radiation.” Mol Cell Biol 18(11): 6423–6429.

Rossi, M. J., S. F. DiDomenico, M. Patel and A. V. Mazin (2021). “RAD52: Paradigm of Synthetic Lethality and New Developments.” Front Genet 12: 780293.

Rossi, M. J., O. M. Mazina, D. V. Bugreev and A. V. Mazin (2010). “Analyzing the branch migration activities of eukaryotic proteins.” Methods 51(3): 336–346.

Singleton, M. R., L. M. Wentzell, Y. Liu, S. C. West and D. B. Wigley (2002). “Structure of the single-strand annealing domain of human RAD52 protein.” Proc Natl Acad Sci U S A 99(21): 13492–13497.

Skinner, S. P., R. H. Fogh, W. Boucher, T. J. Ragan, L. G. Mureddu and G. W. Vuister (2016). “CcpNmr AnalysisAssign: a flexible platform for integrated NMR analysis.” J Biomol NMR 66(2): 111–124.

Storici, F., K. Bebenek, T. A. Kunkel, D. A. Gordenin and M. A. Resnick (2007). “RNA-templated DNA repair.” Nature 447(7142): 338–341.

Struble, L. R., J. J. Lovelace and G. E. O. Borgstahl (2024). “A glimpse into the hidden world of the flexible C-terminal protein binding domains of human RAD52.” J Struct Biol 216(3): 108115.

Sugiyama, T., J. H. New and S. C. Kowalczykowski (1998). “DNA annealing by RAD52 protein is stimulated by specific interaction with the complex of replication protein A and single-stranded DNA.” Proc Natl Acad Sci U S A 95(11): 6049–6054.

Sung, P. (1997). “Function of yeast Rad52 protein as a mediator between replication protein A and the Rad51 recombinase.” J Biol Chem 272(45): 28194–28197.

Tan, J., M. Duan, T. Yadav, L. Phoon, X. Wang, J. M. Zhang, L. Zou and L. Lan (2020). “An R-loop-initiated CSB-RAD52-POLD3 pathway suppresses ROS-induced telomeric DNA breaks.” Nucleic Acids Res 48(3): 1285–1300.

Waudby, C. A., A. Ramos, L. D. Cabrita and J. Christodoulou (2016). “Two-Dimensional NMR Lineshape Analysis.” Sci Rep 6: 24826.

Wei, L., S. Nakajima, S. Bohm, K. A. Bernstein, Z. Shen, M. Tsang, A. S. Levine and L. Lan (2015). “DNA damage during the G0/G1 phase triggers RNA-templated, Cockayne syndrome B-dependent homologous recombination.” Proc Natl Acad Sci U S A 112(27): E3495–3504.

Welty, S., Y. Teng, Z. Liang, W. Zhao, L. H. Sanders, J. T. Greenamyre, M. E. Rubio, A. Thathiah, R. Kodali, R. Wetzel, A. S. Levine and L. Lan (2018). “RAD52 is required for RNA-templated recombination repair in post-mitotic neurons.” J Biol Chem 293(4): 1353–1362.

Williamson, M. P. (2013). “Using chemical shift perturbation to characterise ligand binding.” Prog Nucl Magn Reson Spectrosc 73: 1–16.

Xu, X., S. Vaithiyalingam, G. G. Glick, D. A. Mordes, W. J. Chazin and D. Cortez (2008). “The basic cleft of RPA70N binds multiple checkpoint proteins, including RAD9, to regulate ATR signaling.” Mol Cell Biol 28(24): 7345–7353.

Yasuhara, T., R. Kato, Y. Hagiwara, B. Shiotani, M. Yamauchi, S. Nakada, A. Shibata and K. Miyagawa (2018). “Human Rad52 Promotes XPG-Mediated R-loop Processing to Initiate Transcription-Associated Homologous Recombination Repair.” Cell 175(2): 558–570 e511.

Ying, J., F. Delaglio, D. A. Torchia and A. Bax (2017). “Sparse multidimensional iterative lineshape-enhanced (SMILE) reconstruction of both non-uniformly sampled and conventional NMR data.” J Biomol NMR 68(2): 101–118.

Zhao, W., C. Wiese, Y. Kwon, R. Hromas and P. Sung (2019). “The BRCA Tumor Suppressor Network in Chromosome Damage Repair by Homologous Recombination.” Annu Rev Biochem 88: 221–245.

