## Supplementary figures S1-S4; Supplementary Tables S1-S2 for "RAD52 and RPA act in a concert promoting inverse RNA strand exchange"

**Affiliations:**
[1]Department of Biochemistry and Structural Biology, University of Texas Health Science Center; San Antonio, TX 78229, USA.

[2]Greehey Children's Cancer Research Institute, The University of Texas Health Science Center at San Antonio

[3]Department of Biochemistry, Vanderbilt University, Nashville, TN 37232, USA

[†]Current Address: Laboratoire de Chimie et Biochimie Pharmacologiques et Toxicologiques, Université Paris Cité, France, 75006, Paris

**Contents:**

Supplementary figures S1-S4
Supplementary tables S1-S2

### Suppl. Figure S1

**A**

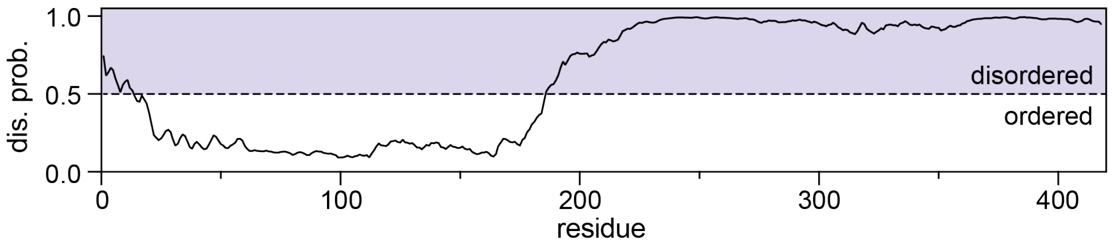


**B**

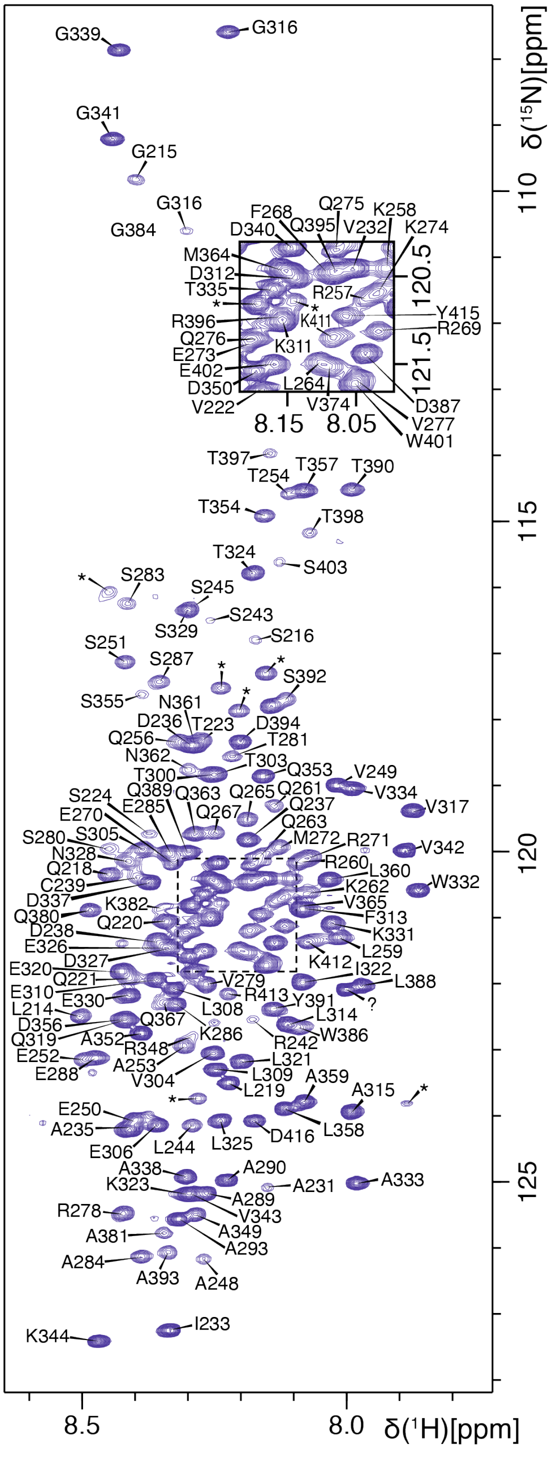

**Figure S1. RAD52 CTD is intrinsically disordered**. (**A**) AIUPred (PMID:38747347) prediction of disorder in full-length RAD51. Residues with values greater than 0.5 are conformationally disordered. (**B**) Assigned $^{1}H$,$^{15}N$-HSQC of RAD52 CTD (217-418), ambiguously assigned and unassigned peaks are denoted with an asterisk.

**Suppl. Figure S2**

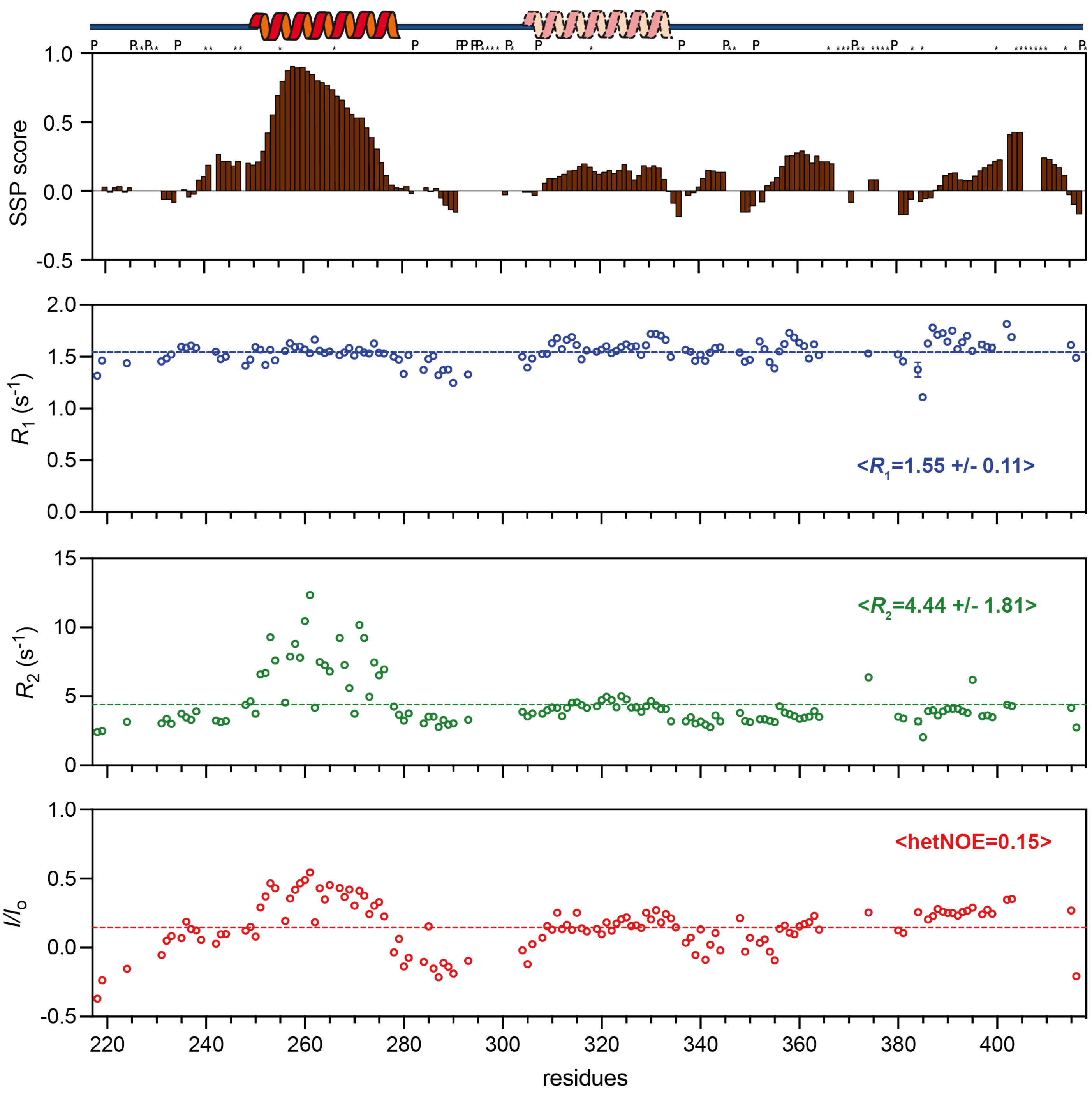

**Figure S2. Secondary structure characteristics of RAD52 (217-418).** (**A**) Secondary structure propensity (SSP) score, (**B**) $R_1$ and (**C**) $R_2$ rates, and the (**D**) $^{1}H$-$^{15}N$ heteronuclear NOE plotted against the RAD52 CTD sequence. The dashed lines represent the average value for each experiment. A cartoon representation of the $\alpha$-helix identified from chemical shift information is shown above the plots. P indicates a proline and asterisks represent unassigned residues.

**Suppl. Figure S3**

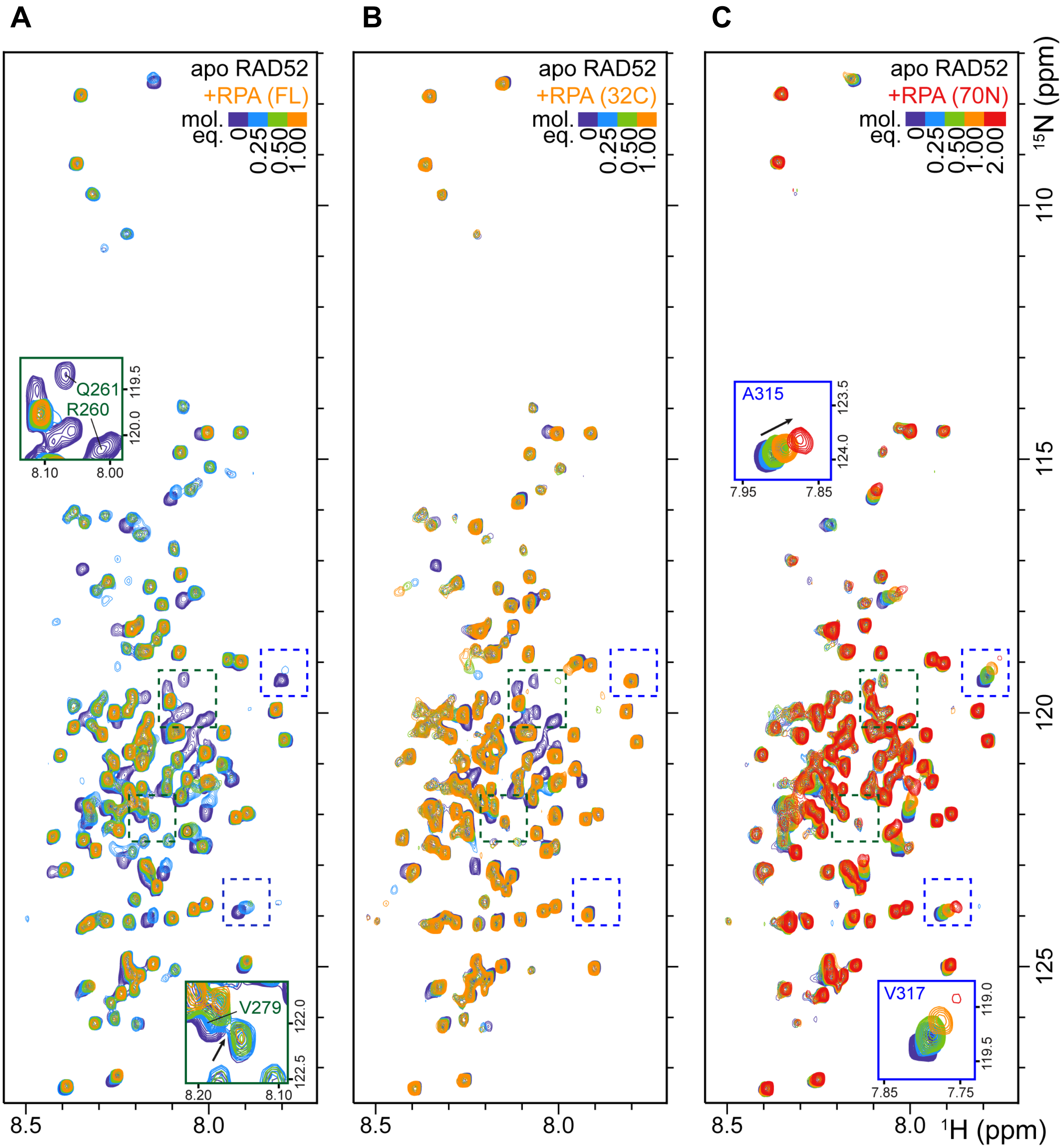

**Figure S3. Representative HSQC spectra for RPA32C and RPA70N titrations.** $^{15}N$, $^{1}H$ HSQC spectra of $^{15}N$ RAD52 CTD (purple) titrated with 0.25 (blue), 0.5 (green), 1 (orange), or 2 (red) molar equivalents of (**A**) RPA full length (FL) heterotrimer, (**B**) RPA32C, or (**C**) RPA70N domains.

#### Suppl. Figure S4

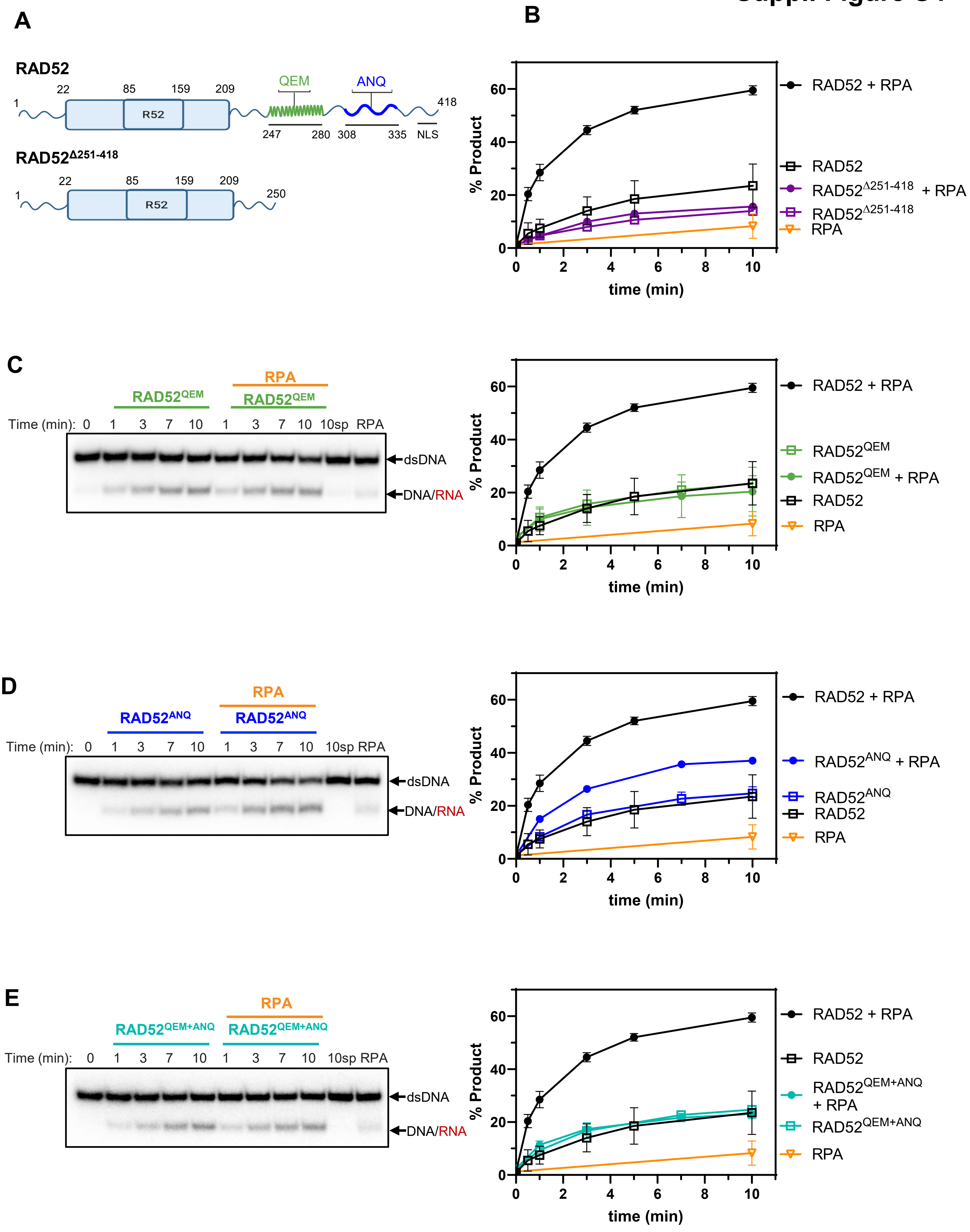

**Figure S4. The effect of RPA on the kinetics of inverse RNA strand exchange promoted by RAD52 or RAD52 CTD mutants.** (**A**) Schematic representation of the RAD52 and RAD52 ΔCTD (RAD52$^{\Delta 251\text{-}418}$) maps. The QEM and ANQ mutations located in the primary (247-280, green) and secondary (308-335, blue) RPA-binding sites, respectively, are highlighted. The inverse RNA strand exchange reaction conditions are the same as in Figure 4. (**B**) The kinetics of inverse RNA strand exchange promoted by RAD52$^{\Delta 251\text{-}418}$ (2 μM) with and without RPA. (**C-D**) Representative polyacrylamide gels, and quantification of inverse RNA strand exchange kinetics promoted by (**C**) RAD52$^{QEM}$ (2 μM), (**D**) RAD52$^{ANQ}$ (1.7 μM), and (**E**) RAD52$^{QEM+ANQ}$ (1.5 μM) with and without RPA. The experiments were repeated at least 3 times; error bars indicate SD.

**Supplementary Table S1. Sequences of the Oligonucleotides used in this Study.**

| Number (no.) | Length (nt) | DNA or RNA | Sequence (5'→3') |
|---|---|---|---|
| **117** | 94 | DNA | TCCTTTTGATAAGAGGTCATTTTTGCGGATGG<br>CTTAGAGCTTAATTGCTGAATCTGGTGCTGTA<br>GGTCAACATGTTGTAAATATGCAGCTAAAG |
| **211** | 48 | DNA | GAAGCATTTATCAGGGTTATTGTCTCATGAGC<br>GGATACATATTTGAAT |
| **1** | 63 | RNA | ACAGCACCAGATTCAGCAATTAAGCTCTAAGC<br>CATCCGCAAAAATGACCTCTTATCAAAAGGA |
| **517** | 63 | RNA | UCCUUUUGAUAAGAGGUCAUUUUUGCGGAUGG<br>CUUAGAGCUUAAUUGCUGAAUCUGGUGCUGU |
| **557** | 63 | RNA | CUGGUGAAAGUAAAAGAUGCUGAAGAUCAGUU<br>GGGUGCACGAGUGGGUUACAUCGAACUGGAU |

**Supplementary Table S2. Protein Expression Vectors used in this Study.**

| Construct | Mutation | Vector backbone | Source |
|---|---|---|---|
| **GST-RAD52-CTD** | Deleted aa 1-216 | pGEX-6P-2 | this study |
| **GST-RAD52-CTD-QEM** | Deleted aa 1-216 point mutants R260Q, Q261E, K262M | pGEX-6P-2 | this study |
| **GST-RAD52-CTD-ANQQAA** | Deleted aa 1-216 Point mutants L314A, V317N, L321Q, I322Q, L325A, W332A | pGEX-6P-2 | this study |
| **GST-RAD52-del251-418** | Deleted aa 251-418 | pGEX-6P-2 | this study |
| **his-RAD52-NTD 1-209** | Deleted aa 210-418 | pET-15b | (Hanamshet and Mazin 2020) |
| **GST-RAD52-ANQQAA** | Point mutants L314A, V317N, L321Q, I322Q, L325A, W332A | pGEX-6P-2 | this study |
| **GST-RAD52-QEM/ANQQAA** | Point mutants R260Q, Q261E, K262M, L314A, V317N, L321Q, I322Q, L325A, W332A | pGEX-6P-2 | this study |
| **RPA** | none | pET11d | (Henricksen, Umbricht, and Wold 1994) |
| **RPA32C** | RPA aa 202-270 | pET-15b | (Mer et al. 2000) |
| **RPA70N-E7R** | RPA aa 1-120, point mutation E7R | pET-15b | (Feldkamp et al. 2013) |
